# Better together: Microbial diversity might facilitate the invasion success of the seagrass *Halophila stipulacea* in mixed Mediterranean seagrass communities

**DOI:** 10.64898/2026.02.09.704841

**Authors:** Emma Hoza-Frederick, Sergio Martínez-Campos, Paul H. Barber, Marlen I. Vasquez, Vasileios Fotopoulos, Chrystalla Antoniou, Katerina Drakou, Filipa Godoy-Vitorino, Kelcie L. Chiquillo

**Affiliations:** Department of Biology, University of Puerto Rico Río Piedras, P.O. Box 23360, San Juan, PR 00931 USA; BETA Technological Centre- University of Vic- Central University of Catalunya (BETA-UVIC- UCC), Carretera de Roda 70, 08500, Vic, Barcelona, Spain; Antimicrobial Resistance Unit, Animal Health Department, Faculty of Veterinary Medicine, Complutense University of Madrid, Madrid, Spain; Department of Ecology and Evolutionary Biology, University of California, Los Angeles, Los Angeles, CA 90095 USA; Department of Chemical Engineering, Cyprus University of Technology, Limassol 3036, Cyprus; Department of Agricultural Sciences, Biotechnology and Food Science, Cyprus University of Technology, Limassol 3036, Cyprus; Department of Microbiology and Immunology, University of Puerto Rico School of Medicine, San Juan, PR 00921 USA

**Keywords:** seagrass, *Halophila stipulacea*, *Cymodocea nodosa*, microbes, 16S rRNA, invasion, Mediterranean, competition, seagrass

## Abstract

Microorganisms are increasingly recognized as key facilitators of invasion success for introduced species into new environments. The globally invasive seagrass *Halophila stipulacea* flourishes in mixed environments with native seagrasses, where it exhibits enhanced growth, while, in contrast, native seagrasses in mixed environments experience reduced growth. Here, we hypothesize that microbes may support the success of invasive seagrass in mixed Mediterranean environments. We analyzed 16S rRNA genes to characterize the microbial diversity on the phyllosphere alongside biochemical, morphological, and sediment nutrient measurements of the Mediterranean-native seagrass *Cymodocea nodosa* and the invasive *H. stipulacea* from a controlled mesocosm experiment. Overall, *C. nodosa* in monoculture harbored a microbiome exhibiting higher ASV richness and a distinct community composition than *H. stipulacea*. Variation in bacterial diversity associated with hydrogen peroxide (H_2_O_2_) and internode length suggests that microbial communities of the native seagrass might be shaped by its stress. Conversely, *H. stipulacea’s* microbiome was most abundant in mixed environments, with bacteria significantly reduced in monoculture, and bacterial diversity loosely associated with growth, suggesting that microbes are critical to assisting and possibly facilitating *H. stipulacea* in mixed environments. Overall, our findings suggest that invasive *H. stipulacea* in the Mediterranean Sea are capable of recruiting beneficial bacteria, creating microbial interactions that support its success, and undermining the resilience of native seagrasses in mixed beds. Future work should center on the mechanisms driving *H. stipulacea* bacterial communities and investigating whether *H. stipulacea* actively determines its own microbiome, or whether its microbiome is passively determined by environmental variables.

## 1. Introduction

Microbial communities are increasingly recognized as key players in plant invasion dynamics and competition (Rudgers et al. 2005; Zhang et al. 2010; Policelli et al. 2018; Elsheikh et al. 2021; Dawkins et al. 2022). Essential to all forms of life and vital for providing resources to their hosts (Rout and Callaway 2012; Dawson and Schrama 2016), as well as driving processes that sustain ecosystems and the planet (Policelli et al. 2018; Z. Zhang et al. 2020; Elsheikh et al. 2021; Hesse et al. 2021; Dawkins et al. 2022). Microbes can influence invasion success and plant competition through several key functions, including nutrient acquisition, stress tolerance, pathogen protection and resource availability (Inderjit and van der Putten 2010; Dawson and Schrama 2016; Dawkins et al. 2022). These functions are mediated by mechanisms such as the production of phytotoxins, phytohormones, secondary metabolites, and disruption of native symbioses (Cipollini et al. 2012), and shifts in microbial communities that can suppress the growth of their competitors (Fang et al. 2022, Dar et al. 2024).

Studies of plant-microbe interactions in marine environments are much scarcer than those in terrestrial environments yet suggest that microbes can be important factors in successful species invasions. For example, the invasive alga *Caulerpa taxifolia* had 200% greater biomass when grown with intact sediment microbes compared to sterile sediment microbes, suggesting that microbes facilitate invasive growth success (Gribben et al. 2017). Additionally, Aires et al. (2013) used microbes to track the invasion source of *Caulerpa racemosa*, suggesting that associated microbes may assist its acclimation to novel habitats. However, the properties of these communities can change over the course of an invasion, as Saha et al. (2016) demonstrated that the macroalga *Gracilaria vermiculophylla* becomes vulnerable to epibionts from its original range, while gaining resistance to epibionts in its invaded territory. Increasing evidence demonstrates that seagrasses’ microbial communities—particularly those associated with roots, rhizomes, and leaves—play a pivotal role in determining establishment to environmental changes (Tarquinio et al. 2019).

Like terrestrial plants, microbes confer beneficial advantages to seagrasses, including enhanced nutrient acquisition, protection against pathogens, biofouling, toxins, and oxidative stress; and promotion of plant growth and developmental processes (Ugarelli et al. 2017). Microbial mediation can explain patterns of establishment and spread across seagrasses among the three known invasive seagrass species—*Zostera japonica,* a highly invasive seagrass in the Pacific Northwest whose associated microbes are detoxifying and denitrifying, growth-promoting, nitrogen-fixing, and epibiont-inhibiting (Wojahn et al. 2016; Crump et al. 2018); *Halophila ovalis*, whose associated microbes can fix carbon and nitrogen, cycle sulfur, promote growth and stress tolerance, and degrade C_1_ products like methanol and methane (Yan et al. 2021); and the globally invasive *Halophila stipulacea* (Lipkin 1975, Harrison and Bigley 1982; Short et al. 2010). Originating from the Red Sea and Indian Ocean, *H. stipulacea* is a Lessepsian migrant that colonized the Mediterranean Sea 150 years ago after the opening of the Suez Canal in 1869. First observed in the Mediterranean in 1894 (Willette et al. 2014), *H. stipulacea* is believed to have subsequently been introduced to the Caribbean Sea via boat (Winters et al. 2020), with first sightings in the Caribbean reported in Grenada in 2002 (Willette et al. 2014).

As an opportunistic, phenotypically plastic species, *H. stipulacea’s* ability to tolerate varying light, depth, temperature, salinity, and hydrodynamic regimes (Rotini et al. 2017) has been associated with its high invasion success, although less work has been done examining how *H. stipulacea’s* microbial communities might influence this plasticity. To date, most evidence supports the host generalist hypothesis, which posits that *H. stipulacea’s* ability to adapt to the conditions of its invaded range comes from effectively exploiting the microbial communities of its invaded range (Parker 2001; Aires et al. 2022), which assists its competition against native seagrasses in mixed communities. In syntopic populations in the Mediterranean Sea, Conte et al. (2023) found that *H. stipulacea* exhibited unaffected density and largely unaffected bacterial diversity when growing next to the native seagrass, while the native seagrass exhibited decreased density and bacterial diversity. In the Caribbean Sea, Aires et al (2021) identified bacteria that putatively promoted growth, nutrient acquisition, stress and saline tolerance, detoxification, quorum sensing inhibition, and epiphyte degradation, potentially enabling *H. stipulacea* to thrive in nonnative territories. Further support comes from Winters et al. (2023), who found that Caribbean *H. stipulacea* exhibited higher growth rates and biomass than native Red and Mediterranean Seas populations, suggesting a shift in *H. stipulacea’s* physiology between native and invaded territories, which could potentially be microbe mediated. Put together, positive impacts on the seagrass holobiont, the relationship between microbes and their seagrass hosts, might enable invaders to be more effective competitors than their rivals by enhancing seagrass ecological functions (Ugarelli et al. 2017; Tarquinio et al. 2019; Conte et al. 2021).

*H. stipulacea* microbial communities may be impacted by the invasive, its environment, or the act of competition itself. It is well understood that plants can mediate their own microbial communities through various genomic, metabolic, and structural strategies (Pascale et al. 2020; Barnes et al. 2025). Cúcio et al. (2016) demonstrated that locally, seagrass-associated microbes are primarily plant-selected, while more broadly, they are determined environmentally. Mejia et al. (2016) observed that *H. stipulacea’s* epiphytic bacterial communities varied along a depth gradient in the Gulf of Aqaba. Similarly, Rotini et al. (2017) demonstrated that abiotic factors such as depth significantly affected *H. stipulacea* microbial community structure. Plant competition has also been shown to alter microbial diversity, composition, and functional potential across multiple systems; for example, competition between native and invasive terrestrial grasses can restructure rhizosphere communities, sometimes amplifying competitive asymmetries (LaForgia et al. 2022; LaForgia et al. 2021), while in crop–weed systems, microbial shifts can mediate allelopathic suppression (Mishra et al. 2013; Kohli et al. 2024). In marine systems, competition between macroalgae (Egan et al. 2017) and coral species (Teplitski et al. 2016) has been linked to changes in surface-associated microbial communities. Taken together, these studies suggest that competition can act as a driver of microbiome change, rather than microbes simply mediating competition.

Despite evidence from controlled mesocosm studies demonstrating that native seagrass facilitates the success of invasive *H. stipulacea* (Chiquillo et al. 2022), the role of microbes in shaping these interactions remains unresolved. Across both the Mediterranean and Caribbean Seas, *H. stipulacea* tends to dominate mixed assemblages, exhibiting enhanced growth when co-occurring with native seagrasses compared to when it grows alone—often to the detriment of native species (Chiquillo et al. 2022). In contrast, native seagrasses *Syringodium filiforme* and *Cymodocea nodosa* lost shoots when grown with *H. stipulacea* at full density and only showed positive growth in monoculture. Invasive species capable of driving their own invasions are typically superior competitors (Fleming and Dibble 2015). However, given the importance of microbes to plants (Barrow et al. 2008), the competitive advantages of *H. stipulacea* may be facilitated by microbes. In this study, we examine to what extent microbial communities contribute to *H. stipulacea* invasion outcomes through (1) *H. stipulacea* actively selecting for microbes that provide a competitive advantage over native seagrasses, (2) stochastic processes shaped by the environment, or (3) *H. stipulacea* competition with native seagrasses altering microbial communities, which it is then able to physiologically benefit from and exploit.

The objective of this study is to determine whether microbial communities differ across competition contexts that are already known to enhance Halophila stipulacea performance in mixed Mediterranean seagrass communities. Using a controlled mesocosm experiment, we compared microbial diversity, community composition, and biochemical indicators in both the invasive *H. stipulacea* and the native *Cymodocea nodosa* under monoculture and mixed conditions. We hypothesized that (1) *H. stipulacea* would harbor more present or beneficial microbes in mixed communities than in monoculture, and (2) microbial assemblages would correlate with physiological traits such as oxidative stress and shoot growth. Our findings offer new insights into marine plant–microbe interactions and the microbial mechanisms that may underlie biological invasions in seagrass ecosystems.

## 2. Materials and Methods

### 2.1. Experimental design of mesocosm

To examine the effects of microbial communities on seagrass species interactions, we expanded upon a previously published experiment (Chiquillo et al. 2022), which manipulated seagrass community composition (monocultures and mixed communities) and density in controlled mesocosms to assess interactions between the native *Cymodocea nodosa* and invasive *Halophila stipulacea*. The experiment included four treatments with ten replicates each: (1) native and invasive grown together at natural densities, (2) native and invasive together at reduced densities, (3) invasive monoculture, and (4) native monoculture. Experimental units were maintained in flow-through mesocosms for six weeks under ambient conditions, from October 16 to December 3, 2018 at the Marine Aquaculture Center within the Fisheries Aquatic Research Center, Larnaka, Cyprus. In this study, we investigate the microbial diversity and composition of the phyllosphere and seagrass tissue from each of these experimental treatments, to test whether seagrass community structure and density impacts associated microbial communities.

### 2.2 Plant descriptors

To evaluate how plant physiological responses shift in response to seagrass competition, as described in Chiquillo et al. (2022), samples were collected from both *H. stipulacea* and *C. nodosa* at the end of the mesocosm experiment to record the biochemical, morphological (e.g., number of shoots), and soil variables. We measured photosynthetic leaf pigments, indirectly linked to photosynthetic activity in each treatment, by extracting photosynthetic pigments (chlorophyll a and chlorophyll b, total carotenoids) from 50mg of blade tissue from at least 10 randomly selected leaves following modified methods of Richardson et al. 2002. Specifically, we determined chlorophyll a and b concentrations (mg/g of fresh weight, FW) from lyophilized samples after extraction with DMSO, recording average absorbances (A) at 470, 652, 665, and 750 nm from duplicate readings on a Spectroquant Pharo 100 spectrophotometer. Carotenoids and chlorophyll concentrations were estimated using equations proposed by Sims and Camon 2002.

To investigate if competition leads to oxidative stress, we measured lipid peroxidation as an indicator of cellular damage. We tested for both hydrogen peroxide (H_2_O_2_) and malondialdehyde (MDA) using five haphazardly selected blades from the replicates within each treatment; these blades were pooled to create three composite samples per treatment. Lipid peroxidation was determined from the measurement of malondialdehyde (MDA) content resulting from the thiobarbituric acid (TBA) reaction (Minotti and Aust, 1987) using an extinction coefficient of 155 mM-1 cm-1. Hydrogen peroxide was quantified using the KI method, as described by Velikova et al. (2000).

Next, to determine whether growing conditions impacted seagrass growth, we measured growth by calculating the difference in the number of shoots (final (F) minus initial (I)) over the 6-week experiment from Chiquillo et al. (2022), as well as above and belowground biomass and internode length taken from cores at the end of experiment.

To determine nutrient levels in the sediment, we cored 50 mL falcon tubes in random locations within each replicate of each mesocosm and combined them to create a composite sediment sample for each treatment before and after the experiment. Additionally, to measure dissolved nutrients in the water, we collected five water samples from each mesocosm before and after the experiment. We filtered water using a vacuum pump and using 0.2uM pore size. We used Phosphate Test kit (1.14848.0002), Nitrite test kit (#1.14776.002), and Nitrate test in seawater (kit #1.14942.001). Extracts were read twice using Spectroquant Pharo 100 spectrophotometer.

### 2.3 Bacterial communities: DNA extraction and 16S rRNA amplicon sequencing of seagrass-associated bacterial community

To investigate how microbial communities might change based on its response to intraspecific seagrass competition conducted by Chiquillo et al. (2022), we sampled seagrass phyllosphere at two time points: the onset and conclusion of the experiment. At the start of the experiment, we randomly collected one blade per species from each of the 10 experimental units, and pooled blades together into four composite samples per treatment, except for the full mixed-density treatment (where both native and invasive grew together in natural density); in this treatment only sediment was disturbed to mimic the disturbance of the other treatments, but blades were not removed (n=16). At the end of the experiment, we again collected one blade of each experimental unit for each species from each treatment and grouped blades together into four composite samples per treatment (n=24). In total, we obtained 40 composite samples for downstream analysis (Figure 1).

**Figure 1.**
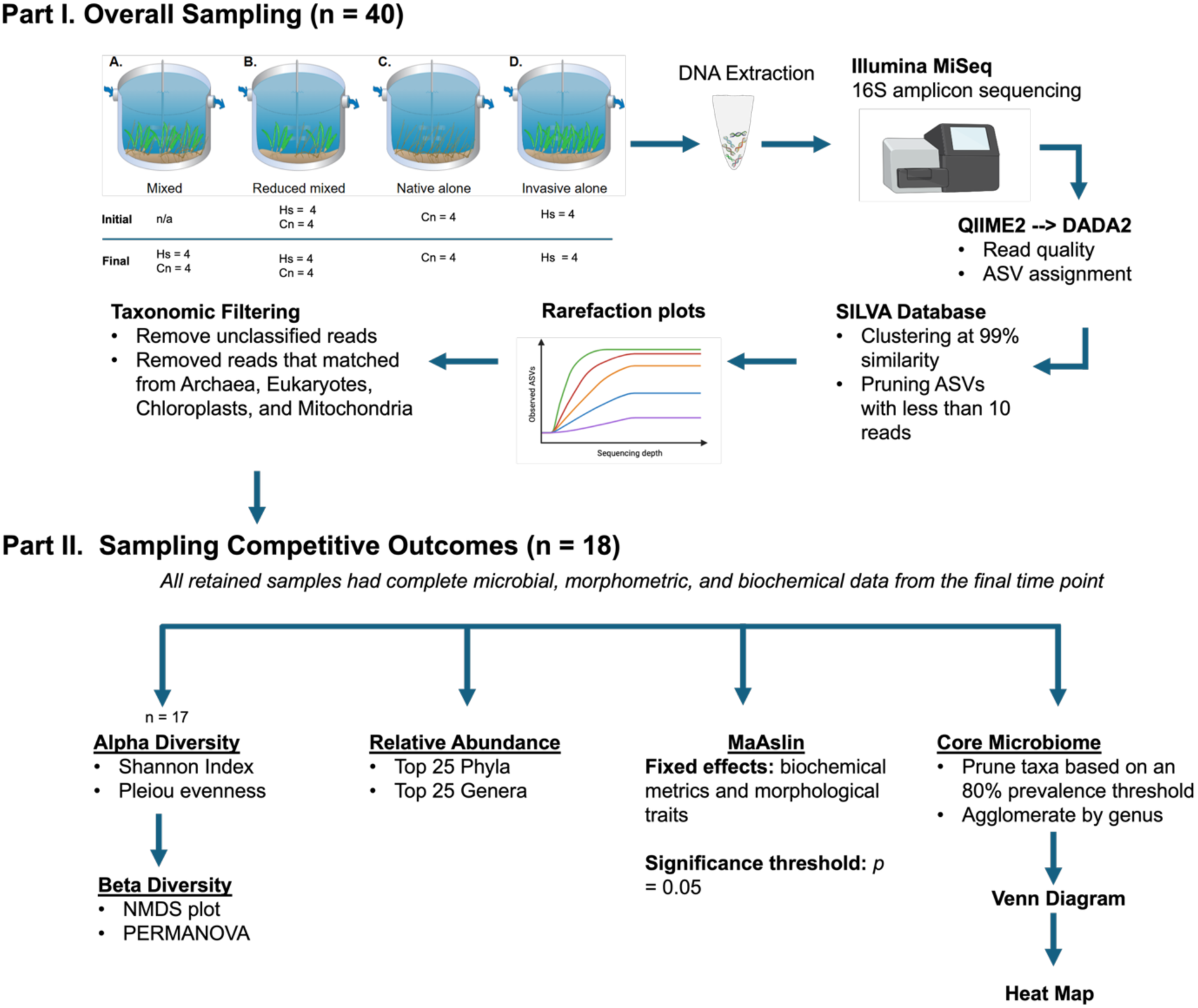
Flowchart of the sampling design and microbial sequencing workflow. Part I – Overall sampling (n = 40). Four mesocosm treatments were established following Chiquillo *et al*. (2022): (A) *Mixed* (native and invasive seagrass species co-occurring at natural densities), (B) *Reduced mixed* (native and invasive species co-occurring at reduced densities to assess density-dependent effects), (C) *Native alone* (monoculture of *Cymodocea nodosa*), and (D) *Invasive alone* (monoculture of *Halophila stipulacea*). DNA extractions were conducted on four composite samples per treatment × species at the initial timepoint (n = 16) and final (n = 24) time points. The full mixed treatment lacked initial samples. Raw sequence reads underwent quality filtering, denoising, chimera removal, and ASV inference using the SILVA database. Sequencing depth adequacy and sample coverage were evaluated using rarefaction curves, maintaining only bacterial taxa. Part II – Sampling Competitive Outcomes (n = 18). Downstream analyses assessed alpha diversity, beta diversity, and differential abundance for taxa under competitive interactions.

To get a bacterial pellet, each sample was scraped from all the epibionts (including small pieces of tissue blade) used for DNA extraction. Tissue blades were gently scraped into sterile Petri dishes using forceps and sterile razors. Each sample was first washed with 10 ml of washing solution (200 mM Tris–HCl pH 8, 10 mM EDTA, and 0.24% Triton X-100; Kadivar and Stapleton 2003). During the scraping process, some tissue fragments were also released into the wash. The resulting supernatant was vortexed (3× 30 s) to detach and collect the bacteria. The plant parts were then removed with sterile tweezers and the remaining solution was centrifuged at 5000 rpm for 20 min to obtain the bacterial pellet for subsequent DNA extraction using the Qiagen Power Soil® DNA isolation kit (Qiagen, Carlsbad, CA, USA), following manufacturer’s instructions.

To characterize the diversity and relative abundance of prokaryotes in the different mesocosm treatments, we sequenced 16S rRNA V4 using primers 515F (GTGYCAGCMGCCGCGGTAA) and 806 R (GGACTACNVGGGTWTCTAAT) (Caporaso et al. 2011) as per the Earth Microbiome Project protocol https://earthmicrobiome.org/protocols-and-standards/16s/. Each sample was amplified in triplicate 25 uL reactions, using the Qiagen Multiplex PCR kit. Thermocycling parameters began with a denaturation step of 94° C for 3 min, followed by 35 cycles of 94° C for 45 sec, annealing for 50° C for 60 sec and extending at 72° C for 90 sec, with a final extension of 72° C for 10 min. Once validated on an agarose gel, triplicate reactions were pooled and cleaned using Agencourt AMPure magnetic beads. Further, we evaluated taxonomic composition with morphological, biochemical traits and soil nutrient (nitrate/nitrite) assays, allowing us to identify microbial indicators associated with competitive outcomes.

### 2.4 Sequencing and Bioinformatics

Pooled PCR samples were dual-indexed using the Illumina Nextera XT Indexing kit and then cleaned a second time with AMPure magnetic beads. Indexed samples were then quantified using a Qubit dsDNA BR kit from Invitrogen to ensure that samples were pooled into equimolar ratios. The pooled library was subsequently sent for sequencing on an Ilumina Miseq v3 with a 20% PhiX at the TCGB core at UCLA (300 bp x 2).

To conduct quality control, amplicon sequence variant (ASV) taxonomy assignment, and community diversity analyses, we processed sequencing data using the QIIME 2 (v. 2024.5.0) microbiome data science platform (Bolyen et al. 2018). Briefly, we demultiplexed and denoised raw sequence data using the DADA2 pipeline (Callahan et al. 2016) and merged the output into a feature table for analysis. Sequences were then trimmed at 10 bp and truncated at 270 and 215 bp for the forward and reverse directions, respectively. Next, ASVs were assigned taxonomy using a naive Bayes taxonomy classifier trained on the SILVA bacterial database (Quast et al. 2013), employing clustering at 99% similarity, and pruning ASVs with less than 10 reads. Rarefaction curves were used to determine whether we measure sufficient sequencing depth given the diversity within the sample. We removed reads that matched archaea, chloroplasts, eukaryotes, and mitochondria to ensure we analyzed only reads belonging to bacteria. Subsequent analyses were conducted in R (v. 4.4.1) (R Core Team 2024), using the ‘qiime2R’ package to translate QIIME 2 artifacts (Bisanz 2025).

16S rRNA microbial compositions of the phyllosphere were analyzed for both native and invasive seagrass species. Using observed ASVs, we calculated Shannon diversity and Pielou evenness with the ‘otuSummary’ package (Yang 2023). Overall diversity between groups (beta diversity) was calculated using a non-parametric Kruskal-Wallis test (pairwise). Additionally, we used a linear model to assess treatment effects on beta diversity metrics, alongside nonparametric tests. To assess community composition of our samples, we produced a non-metric multidimensional scaling (NMDS) plot with Bray-Curtis distances using the ‘vegan’ package in R (Oksanen et al. 2025).

We created a principal coordinate analysis (PCoA) plot to explore how environmental factors, such as our plant descriptors, influenced bacterial compositions in our samples using the full n=18 samples. The plot and covariate statistics were produced using ‘vegan’, ‘ecodist’, and 999 permutations (Oksanen et al. 2025; Goslee and Urban 2007), after which we ran PERMANOVA and ANOSIM tests to detect overall group differences and a pairwise PERMANOVA test to detect pairwise differences with ‘pairwiseAdonis’ (Martinez Arbizu 2025) with 999 permutations. Core microbiomes were determined in R as taxa appearing within at least 80% of samples as done in Aires et al. (2021). Venn diagrams comparing the core microbiomes of our samples were generated using the ‘venn’ package (Dusa et al. 2004). Heatmaps were then produced using ‘pheatmap’ (Kolde 2009) to determine how samples clustered and which taxa within the core microbiomes were mostly present.

To visualize bacterial compositions of our samples, we created bar plots of relative taxonomic abundance using ‘ggplot2’ (Wickham et al. 2025). To identify and test for significant associations of microbial taxa with experimental treatments, we used Microbiome Multivariable Associations with Linear Models (MaAsLin 2; Mallick et al. 2021), an R package designed to assess multivariable associations by integrating microbial data with a broad spectrum of experimental factors correcting for multiple comparisons. In this study, the fixed effects incorporated into the model include biochemical analyses, morphological characters, with taxon relative abundance modeled as a function of treatment and associated metatdata. Multiple testing correction was performed using the false discovery rate (Bejamani-Hochberg), and association were considered significant at a q-values of 0.05. Prior to model fitting, metadata variables were evaluated for collinearty and alignment with the experimental design to ensure that no predictors (i.e. number of shoots) were downstream consequences of treatment effect. Identified associations reflect taxonomic patterns, and any functional roles inferred from these results are framed as hypotheses rather than direct functional evidence.

## 3. Results

### 3.1 Plant descriptors

Native *C. nodosa* had higher photosynthetic pigment levels, including chlorophyll a, chlorophyll b and carotenoids, when grown in either ambient or reduced mixed communities than when grown alone (Table 1). This pattern was opposite for the invasive *H. stipulacea*, resulting in a significant interaction (p < 0.0001 for chlorophyll a, chlorophyll b and carotenoids; Supplemental Table S1). Similarly, there was a significant difference in hydrogen peroxide (H_2_O_2_), a marker of cellular stress, between the two species of seagrass (one way ANOVA; df=12; p < 0.0001; Supplemental Table S1), with H_2_O_2_ concentrations of the native species (mean = 4.18 ± 0.42 SE µmol/g) being more than a seven fold distance than that of the invasive (mean = 0.62 ± 0.09 SE µmol/g), with only a significant interaction between species (p < 0.0001; Supplementary Table S1). Similarly, the native seagrass *C. nodosa* had higher malondialdehyde (MDA) levels than the invasive seagrass (LME; df=12, p <0.05, Supplemental Table S1). Average MDA levels in native *C. nodosa* were 9.18 ± 0.84µmol/g, and for invasive *H. stipulacea* was 7.83 ± 0.34 µmol/g.

**Table 1.** Averages of biochemical, morphological, and soil parameters measured across native and invasive seagrass species under six treatments: Native Mixed, Native Reduced Mixed, Native Alone, Invasive Mixed, Invasive Reduced Mixed, and Invasive Alone, and the standard error. Biochemical parameters include chlorophyll *a* (Chl a), chlorophyll *b* (Chl b), total chlorophyll (Total Chl), chlorophyll *a/b* ratio (Chl a/Chl b), carotenoids (Car), Lipid peroxidation malondialdehyde (MDA), and hydrogen peroxide (H_2_O_2_). Morphological traits include the number of shoots and internode length. Soil parameters include nitrate and nitrite concentrations. Asterisks indicate significant effects from the linear mixed effects model: * p < 0.001for the interaction Treatment × Species. Bolded parameters are values analyzed and shown in Supplemental Table 1.

| Parameter | Native mixed | Native reduced mixed | Native alone | Invasive mixed | Invasive reduced mixed | Invasive alone |
| --- | --- | --- | --- | --- | --- | --- |
| **Chl a\***<br>(mg/FW) | 0.006 ± 0.002 | 0.007 ± 0.000 | 0.005 ± 0.001 | 0.005 ± 0.001 | 0.003 ± 0.001 | 0.004 ± 0.001 |
| **Chl b \***<br>(mg/FW) | 0.003 ± 0.001 | 0.004 ± 0.002 | 0.003 ± 0.001 | 0.003 ± 0.000 | 0.002 ± 0.001 | 0.002 ± 0.001 |
| **Chl a / Chl b** | 2.11 ± 0.84 | 1.75 ± 0.58 | 1.75 ± 0.43 | 1.56 ± 0.19 | 1.56 ± 0.77 | 1.89 ± 0.54 |
| **Car\*** | 0.002 ± 0.001 | 0.002 ± 0.000 | 0.001 ± 0.001 | 0.002 ± 0.001 | 0.001 ± 0.000 | 0.001 ± 0.001 |
| **MDA\***<br>(μmol/g) | 9.33 ± 1.76 | 7.67 ± 0.49 | 10.56 ± 1.79 | 7.27 ± 0.35 | 8.45 ± 1.22 | 7.78 ± 0.18 |
| **H<sub>2</sub>O<sub>2</sub>\***<br>(μmol/g) | 3.73 ± 1.05 | 3.80 ± 0.42 | 5.01 ± 0.90 | 0.49 ± 0.26 | 0.59 ± 0.20 | 0.78 ± 0.06 |
| Change in number of shoots | -1 ± 0.4 | 0.2 ± 0.1 | 1.7 ± .1 | 3.7 ± 1.3 | 4.3 ± 1.5 | 0.5 ± 1.00 |
| Aboveground biomass (g) | 0.18 ± 0.03 | 0.22 ± .04 | 0.55 ± 0.06 | 0.46 ± 0.08 | 0.37 ± 0.08 | 0.38 ± 0.06 |
| Belowground biomass (g) | 1.92 ± 0.35 | 1.71 ± 0.36 | 3.15 ± 0.74 | 2.22 ± 0.39 | 1.79 ± 0.26 | 1.32 ± 0.23 |
| **Internode length\* (cm)** | 3.17 ± 0.46 | 2.78 ± 0.20 | 2.08 ± 0.33 | 0.99 ± 0.04 | 1.05 ± 0.17 | 1.10 ± 0.00 |
| **Nitrates (mg/L)** | 0.93 ± 0.23 | 0.97 ± 0.12 | 1.00 ± 0.17 | 0.93 ± 0.23 | 0.97 ± 0.12 | 1.10 ± 0.00 |
| **Nitrites (mg/L)** | 0.06 ± 0.00 | 0.02 ± 0.00 | 0.05 ± 0.00 | 0.06 ± 0.00 | 0.02 ± 0.00 | 0.06 ± 0.02 |

Average nitrate concentrations in ambient mixed sediment were 0.93 ± 0.23 SE mg/L, while nitrite levels averaged at 0.06 ± 0.04 SE mg/L (Table 1). Similar values were observed in reduced mixed densities, with nitrates at 0.97± 0.12 SE mg/L and nitrites at 0.02 ± 0.03 SE mg/L. Native alone treatments had an average nitrate of 1.00 ± 0.17 SE mg/L and nitrite concentration of 0.053 ± 0.004 SE mg/L respectively, while the invasive showed 1.10 ± 0 SE mg/L for nitrates and 0.06 ± 0.02 SE mg/L for nitrites. Overall, nitrate and nitrite concentrations did not significantly differ between treatments and species.

Morphological values for the number of aboveground and belowground shoots, presented in Table 1 are published in Chiquillo et al. (2022), and no additional statistical analyses were conducted on these data. Briefly, mesocosm experiments in the Caribbean and Mediterranean showed that *H. stipulacea* performed better with native species than alone, while the native species was directly affected by *H. stipulacea*; where it lost shoots and biomass in the presence of the invasive. In the Mediterranean, *H. stipulacea* increased by 3.7 ± 1.3 shoots (+59.2%) when mixed but showed no change alone, whereas the native *Cymodocea nodosa* lost −1 ± 0.4 shoots (−29.2%) when mixed. Aboveground biomass patterns were similar: natives produced significantly more biomass when alone (0.55 ± 0.06 g) than mixed (0.18 ± 0.03 g; p < 0.01), while the invasive species maintained or increased biomass in mixed treatments. In the reduced-density mixed treatment, *H.stipulacea* grew by 4.3 ± 1.5 SE shoots, an increase from an initial of +150.0% ± 50.7 SE (initial shoot density 2.9 ± 0.1). Surprisingly, when density was reduced in the mixed community, the native increased in shoots by 0.2 ± 0.1 SE, which is a +15.0% ± 10.7 SE increase. When *H.stipulacea* grew in reduced mixed treatments, final aboveground biomass was 0.37 ± 0.08 g (estimated initial 0.24 ± 0.03 g), suggesting considerable regrowth during the experiment. When the native seagrass grew in the reduced-density mixed treatment, its aboveground biomass of 0.22 ± 0.04 g was not different from when grown in the ambient density mixed treatment, despite the experimental reduction. However, internode length was not included in Chiquillo et al. 2022, and we found a significant interaction in the length of the rhizomal internodes between species; the average internode length of the native species was 3-fold longer than the invasive (Table 1; Supplementary Table S1).

### 3.2 Native and invasive seagrass phyllosphere

The 16S rRNA raw sequence data have been deposited in the SRA of NCBI with BioProject Number PRJNA1291277 and assigned accession numbers SRR34539037-SRR34539076.

A total of 301,907 reads were sequenced from the original 40 samples. After quality control, joining of paired-end reads, and removal of chimeras, 169,068 high-quality reads remained, representing approximately ∼56% of the original dataset.

Rarefaction analysis was conducted on all 40 samples originally sequenced, demonstrating adequate sequencing depth with a maximum of 46,777 reads, indicating sufficient sampling effort to capture the bacterial diversity present (Supplemental Figure S1A). The remaining treatments—mixed at full density and mixed at reduced density—for each species exhibited intermediate values for both sequencing depth and ASV richness, with no consistent trend across species combinations or density levels.

A total of 18 samples were retained for microbial analyses after completing morphometric, biochemical, and soil nutrient (nitrate/nitrite) assays. To standardize sampling effort for diversity analyses, we rarefied our n=18 samples to a sequencing depth of 1500, which resulted in a loss of one of our samples. Rarefaction curves for 17 samples, which were used exclusively for alpha and beta diversity analyses (Figure 1 Part II), reached saturation at a maximum sequencing depth of 32,142 reads, confirming that sequencing coverage was sufficient to represent the microbial diversity in the dataset (Supplemental Figure S1B). The native-alone and native-reduced treatments exhibited the steepest initial slopes and highest plateau values, consistent with their elevated ASV richness. In contrast, samples from the invasive-mixed and invasive reduced mixed treatments plateaued at lower read depths, reflecting reduced bacterial diversity in those communities.

All alpha and beta diversity calculations were done with rarefied data and using a max of 17 individuals per sample. All remaining analyses, including relative abundance, core microbiome identification, and MaAsLin2 modeling, were performed on the final dataset (n=18), comprising of 826 ASVs, and the average number of ASVs per sample was 220.

### 3.3 Alpha and beta diversity identified of the phyllosphere

Among the 826 bacterial ASVs identified across all taxonomic levels, native and invasive seagrasses portrayed analogous Shannon alpha bacterial diversity measurements (Figure 2A), with no significant relationships between treatment and species (p = 0.171; Supplementary Tables S2), although a linear model found a significant relationship between treatment groups and diversity (intercept p = 2.82e-11). Pielou evenness also showed no significant relationships between groups (p = 0.763; Supplementary Table S3), although a significant linear relationship was detected between treatment group and evenness (intercept p = 7.33e-09).

**Figure 2.**
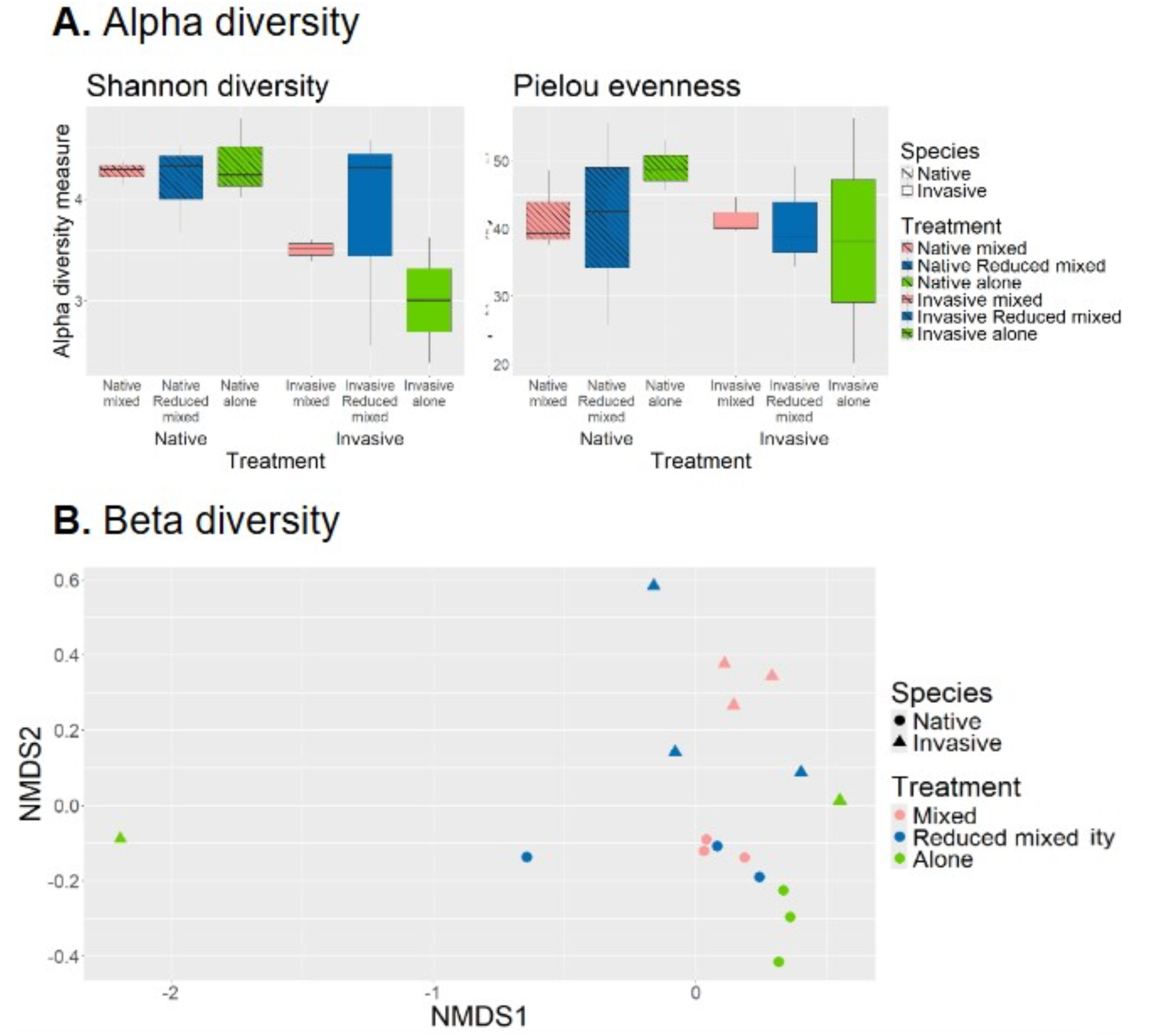
Alpha bacterial diversity of the seagrass samples split by treatment and species. Points are color-coded according to treatment (in ambient fully mixed (pink), reduced mixed (blue) and alone (green). Stippled box plots represent the native *Cymodocea nodosa*, while solid box plots represent the invasive *Halophila stipulacea*. (A.) Shannon (p= 0.171) and Pielou alpha diversity (p=0.763) measures were analyzed on the species level. (B.) Beta bacterial diversity of native versus invasive seagrasses grown in ambient fully mixed (pink), reduced mixed (blue) and alone (green) in a non-metric multidimensional scaling (NMDS) plot with Bray-Curtis distances analyzed on the species level. Circles represent the native *Cymodocea nodosa*, while triangles represent the invasive *Halophila stipulacea*.

Beta diversity analysis using a non-metric multidimensional scaling (NMDS) plot with Bray-Curtis distances found species-level differences between groups overall (PERMANOVA p-value = 0.001; ANOSIM p-value = 0.001; betadisper PERMANOVA p-value: 0.001). Overall, native and invasive seagrass samples displayed a clear separation, where they clustered away from each other (Figure 2B). Yet the pairwise PERMANOVA test found no differences between any two specific groups (Supplementary Table S4).

The top phyla associated with the phyllosphere of native and invasive seagrass showed *Proteobacteria* (Figure 3A), which accounted for 72.1% of the analyzed sequence reads across all samples. The second and third most abundant phyla across all samples were *Bacteroidota* (7.7%) and *Planctomycetota* (3.7%). Additional common phyla found among all groups were *Actinobacteriota* (2.7%), *Verrucomicrobiota* (2.5%), *Desulfobacterota* (2.4%), and *Myxococcota* (1.6%).

**Figure 3.**
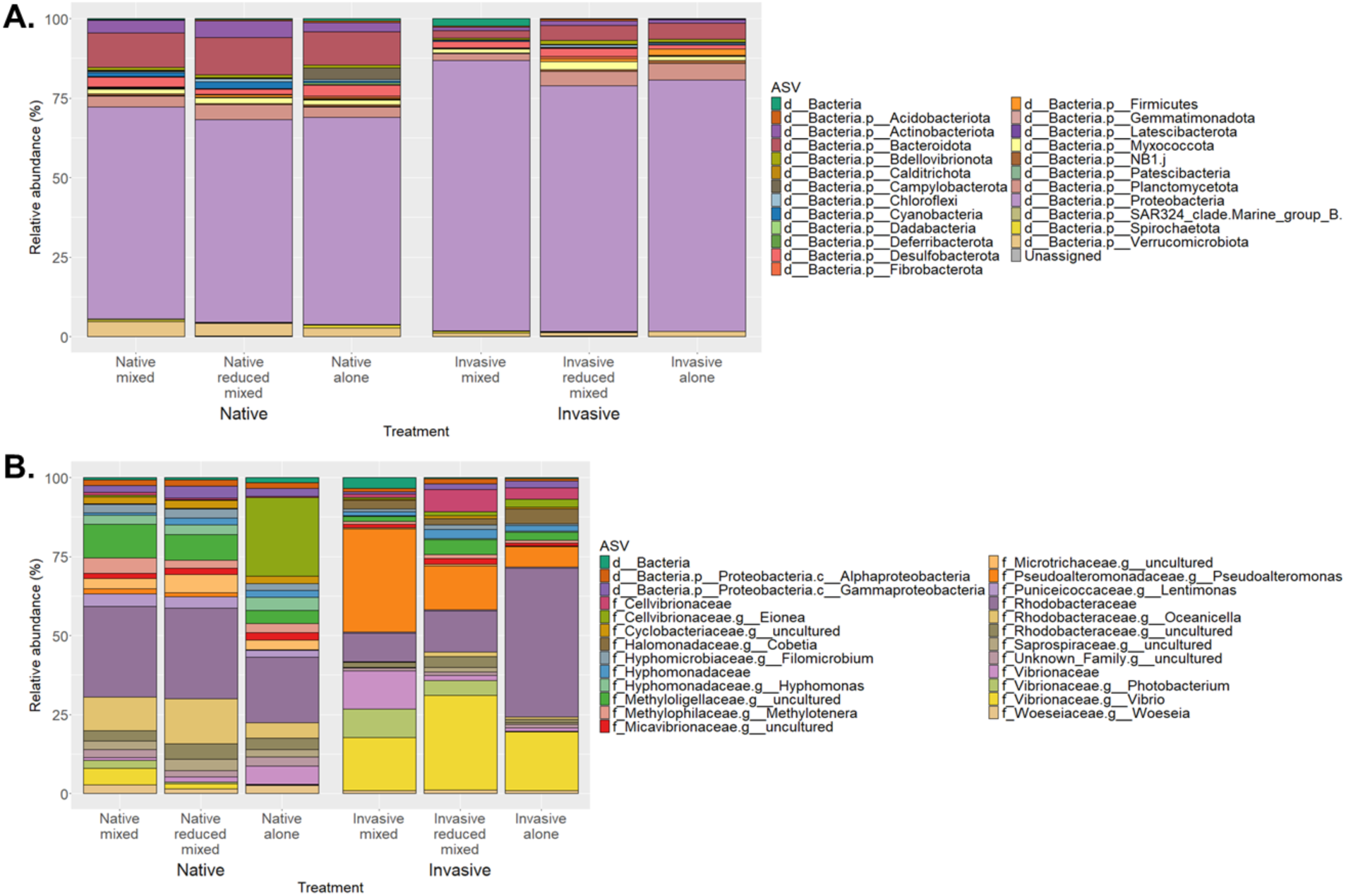
Bacterial relative abundance plots of the seagrass samples split by species and treatment. (A.) Relative abundances of the top 25 most common phyla in native and invasive seagrass samples. (B.) Relative abundances of the top 25 most common genera observed in native and invasive seagrass samples.

The top 25 most abundant genera marked strong differences in taxonomic proportions between the two species of seagrass (Figure 3B). The top genus detected was *Rhodobacteraceae*, which accounted for 26.1% of the analyzed sequence reads across all samples. Surprisingly, the native seagrass *C. nodosa* showed relatively elevated levels of *Bacteroidota* (10.5-11.8%) compared with the invasive seagrass (2.3-5%). We also found higher proportions of uncultured genera of the family *Oceanicella* (5-14.2%)*, Microtrichaceae* (3.1-5.8%), and other genera such as *Hyphomonas* (3-4.3%), *Methylotenera* (2.5-4.9%), and *Lentimonas* (2.1-3.9%) in the native seagrass compared to the invasive seagrass, where it was absent or low in recognizable reads. Compared to the native seagrass, which were absent or low in recognizable reads, there was an increased abundance of the genus *Cobetia* in the invasive *H. stipulacea* (2.1-4.6%).

### 3.4 Bacteria significantly associated with Cymodocea nodosa grown in mixed densities versus alone

Of the top 25 taxa, a family within the *Rhodobacteraceae* in the native seagrass *C. nodosa* possessed a higher relative abundance of the genus *Oceanicella* when grown in mixed density (10.6%) and reduced mixed densities (14.2%) compared to when it grew alone (5%). Additionally, we found higher relative abundance of an uncultured taxon of the family *Methyloligellaceae* in mixed (10.6%) and reduced mixed (8.1%) treatments compared to when it grew alone (4.1%). When the native seagrass grew alone, the native seagrass had elevated levels of the genus *Eionea* (24.9%) versus when it grew with the invasive in mixed (0.6%) and reduced mixed habitats (0.1%). Lastly, when native *C. nodosa* grew alone, we identified elevated levels of the phyla *Camplyobacterota* (3.6%) when taxa abundance was very low or unrecognizable in all other treatments.

Results from MaAslin2 analysis showed that when the native *C. nodosa* grew alone there were three significantly positive associations with the genera *Desulfobulbus* (B = 1.107, q-value= 0.00214), *Bermanella* (B = 1.152, q- value= 0.002), and the PS1 clade of the order *Parvibaculales* (B = 3.916, q- value= 0.003) compared to when it grew mixed or in reduced mixed densities (Figure 4A). We found an increased association with the families *Arcobacteraceae* (B = 5.677, q-value = 0.048), *Desulfococcaceae* (B = 1.445, q-value = 0.083), the genera *Sediminispirochaeta* (B = 1.482, q-value = 0.048) and *Aminicenantales* (B = 4.232, q-value = 0.071). In contrast, the genus *Vibrio* was negatively associated with the native *C. nodosa* when it grew alone compared to when it grew in mixed and reduced density communities (B = - 4.530, q-value = 0.019).

**Figure 4.**
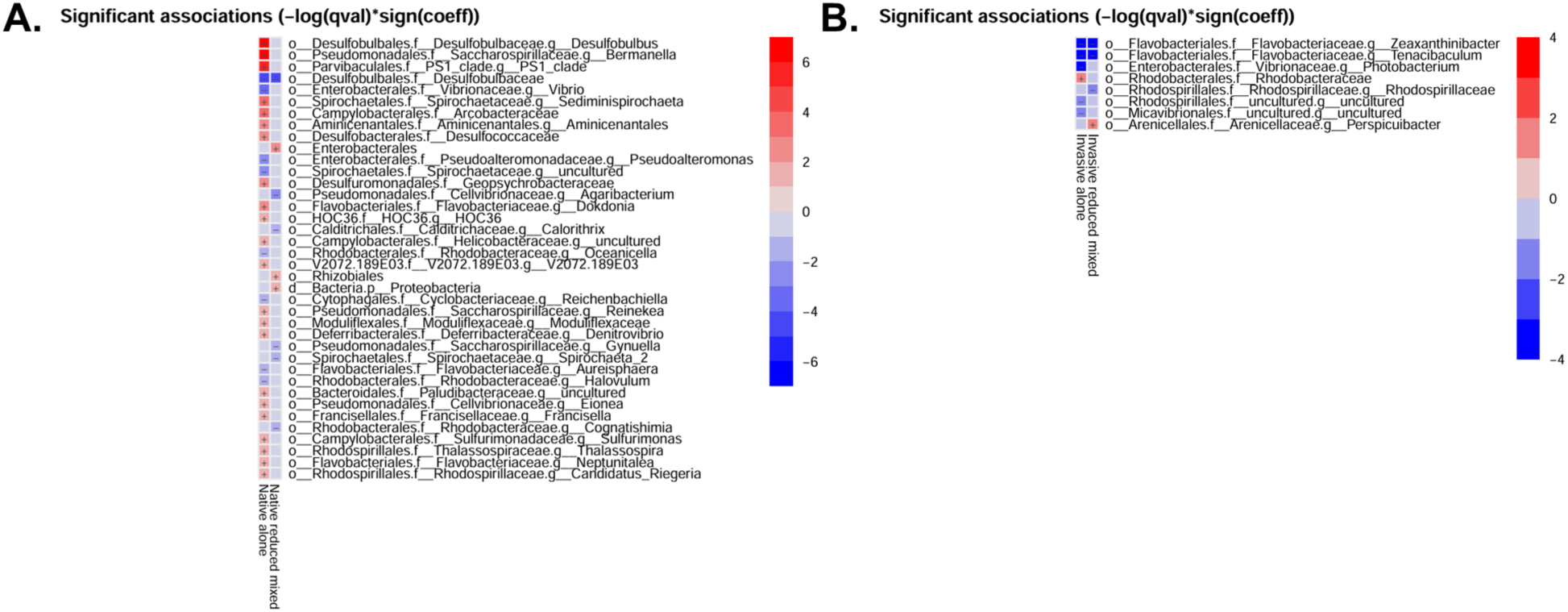
Microbiome Multivariable Associations with Linear Models (MaAsLin 2) heatmaps. A red color (+) indicates a strong positive correlation, while a blue color (-) indicates a strong negative correlation. P-value cutoff value was 0.05. A). Heatmap of taxa correlated with significant changes in native groups only compared to the native mixed group with low number of shoots. B). Heatmap of taxa correlated with significant changes in invasive groups only compared to the invasive mixed group with a high number of shoots.

### 3.5 Bacteria significantly associated with the invasive Halophila stipulacea grown mixed versus alone

The top 25 taxa within the invasive *H. stipulacea* samples were generally dominated by the family *Pseduoaltermonadaceae* and *Vibrionaceae.* Within the family *Pseduoaltermonadaceae*, the genus *Pseudoalteromonas* was notably pronounced across the invasive seagrass treatments. However, when *H. stipulacea* grew in mixed densities, results showed a higher relative abundance of the genus *Pseudoalteromonas* (32.7%) compared to when it grew in a reduced mixed community (14%) and when it grew alone (6.5%). Specifically, *H. stipulacea* showed a pronounced presence of the genus *Vibrio* and *Photobacterium* in mixed density (16.7% and 9.1%, respectively) and reduced mixed density treatments (30% and 4.7%, respectively) versus when it grew alone (18.6 and 0.1%, respectively). Additionally, we found *H. stipulacea* had higher relative abundance of unassigned genera of the family *Cellvibrionaceae* in reduced mixed densities (7%) compared to when it grew in mixed densities (1.1%) and when it grew alone (3.6%).

Results from MaAsLin2 analyses showed that when invasive *H. stipulacea* seagrass grew alone and in reduced densities there was a negative association the genera *Zeaxanthinibacter* (B = -1.175, q-value= 0.015) and *Tenacibaculum* (B = -1.348, q-value= 0.027) compared to when it grows in mixed densities (Figure 4B). Interestingly, the genus *Photobacterium* (B = -6.877, q-value= 0.036) was negatively associated with the invasive *H. stipulacea* when it grew alone compared to when it grew in mixed and reduced density communities. The family *Rhodobacteraceae* was positively associated with the invasive seagrass when it grew in monoculture compared to when it grew in mixed environments (B = 2.478, q-value = 0.139). When the invasive *H. stipulacea* was grown in reduced density there was a negative association with the genus *Rhodospirillaceae* (B = -1.699, q-value= 0.152) and a positive association with the genus *Perspicuibacter* (B = 3.008, q-value = 0.176) compared to when it grew mixed or in monoculture.

### 3.6 Environmental covariates associated with native and invasive bacterial diversity

The relationship between beta diversity and environmental covariates revealed a clear clustering between native and invasive seagrass samples forming distinct groups (Figure 5). Invasive seagrass displayed a broader ordination space, consistent with higher heterogeneity, suggesting more variable microbial assemblages relative to the native samples. Among the environmental covariates, hydrogen peroxide (H_2_O_2_)—a proxy for cellular stress—was significantly associated with microbial community structure of native *C. nodosa* (p = 0.003, R² = 0.610; Table 2). Additionally, covariate internode length, indicative of space-foraging or habitat-seeking behavior, was also significantly correlated with bacterial composition of native *C. nodosa* (p = 0.008, R² = 0.492). However, the number of shoots, often linked to overall growth and productivity, showed a weaker association with the invasive *H. stipulacea* (p = 0.060, R² = 0.329).

**Figure 5.**
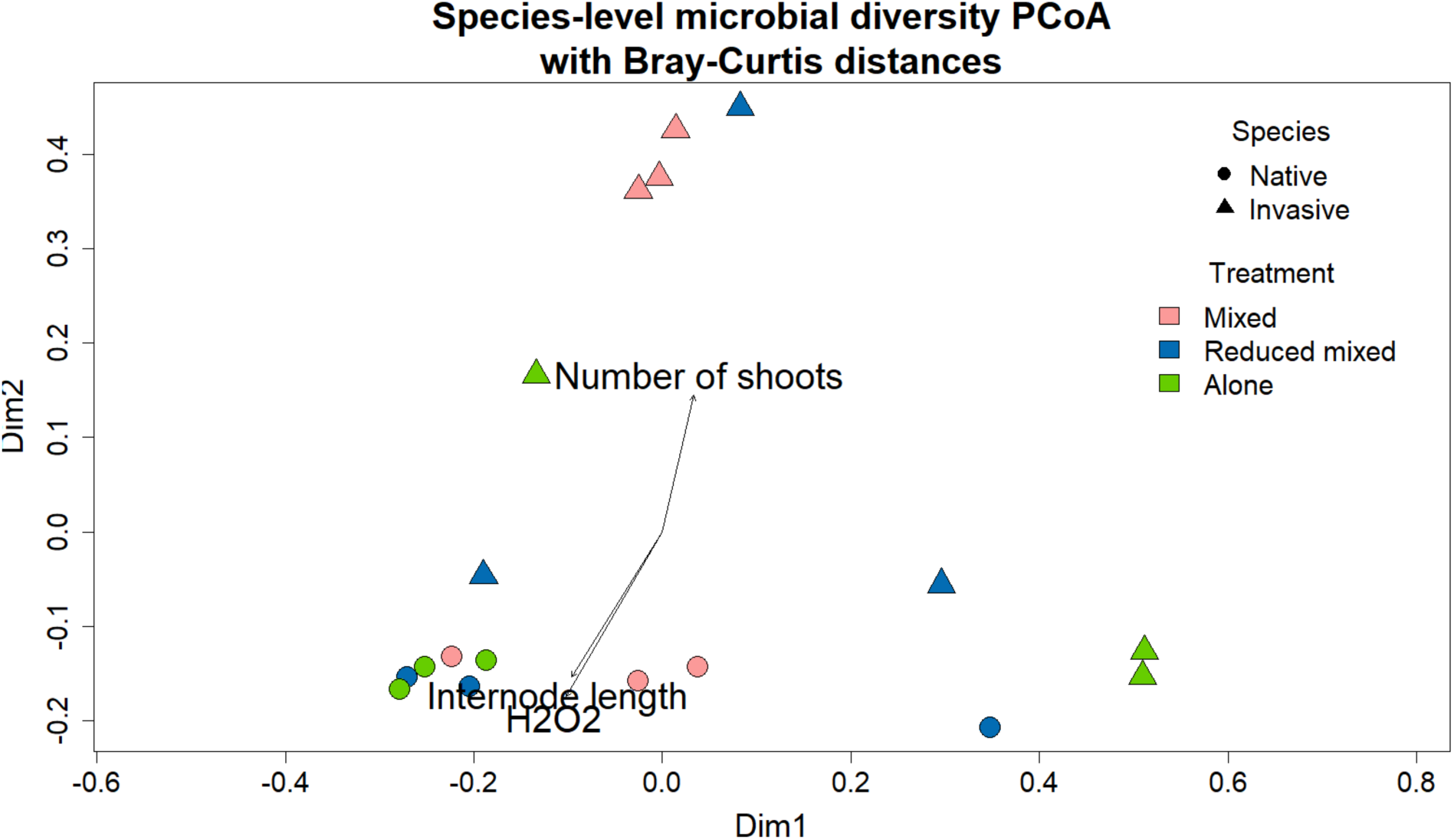
Principal coordinate analysis (PCoA) plot with Bray-Curtis distances on the species level. Native *C. nodosa* samples are mapped as circles, while invasive *H. stipulacea* samples are mapped as triangles. Points are color-coded according to treatment. Arrows signify the strength and direction of correlations between environmental covariates and the ordination plot. Only the top three statistically significant environmental covariates are displayed.

**Table 2.**
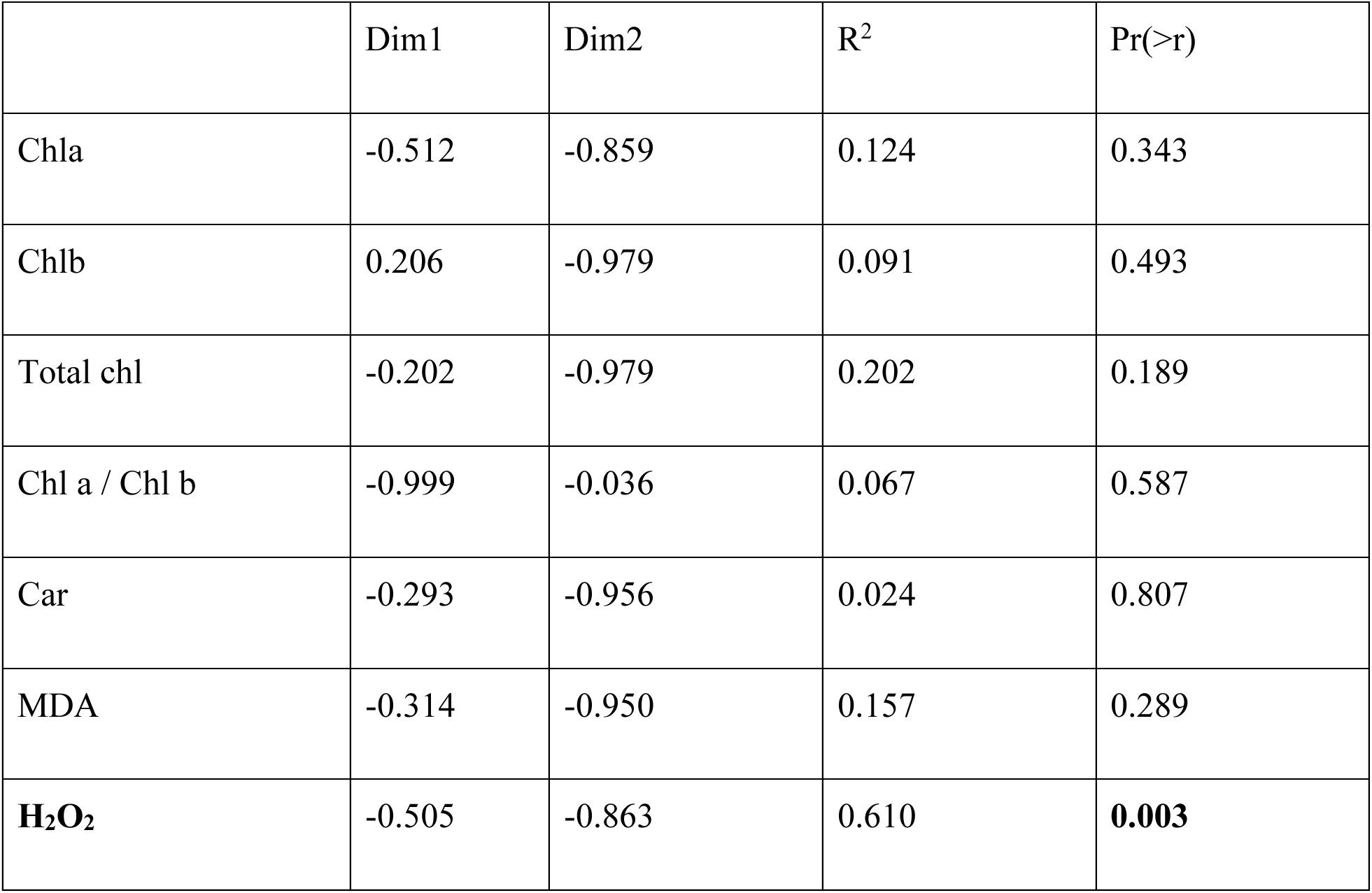
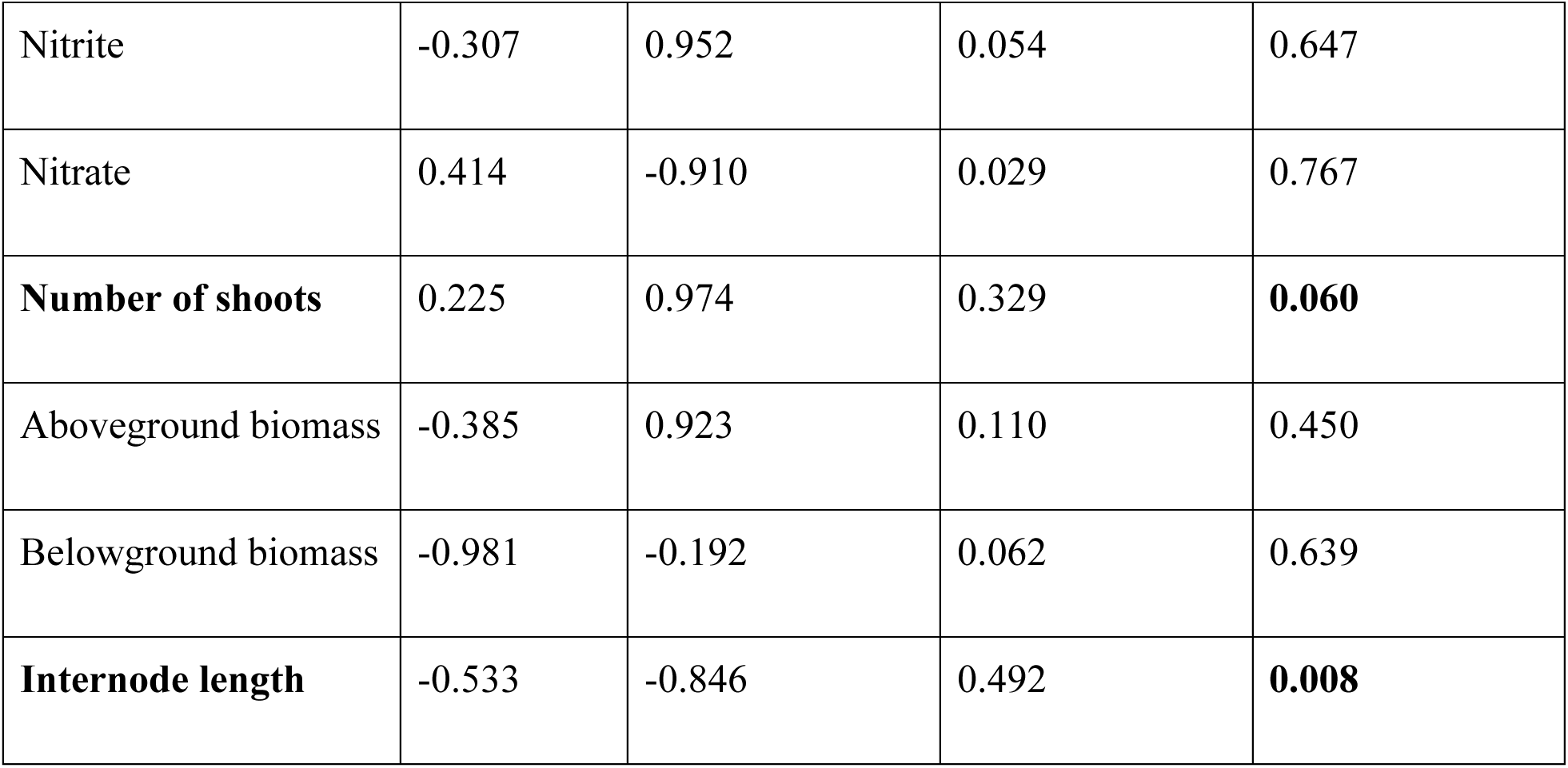
Environmental covariate statistics from PCoA. Significant values are bolded. Chlorophyll a, Chlorophyll b, Carotenoids, MDA and H_2_O_2_. Numerator (numDF) and denominator (denDF)

|  | Dim1 | Dim2 | R<sup>2</sup> | Pr(>r) |
| --- | --- | --- | --- | --- |
| Chla | -0.512 | -0.859 | 0.124 | 0.343 |
| Chlb | 0.206 | -0.979 | 0.091 | 0.493 |
| Total chl | -0.202 | -0.979 | 0.202 | 0.189 |
| Chl a / Chl b | -0.999 | -0.036 | 0.067 | 0.587 |
| Car | -0.293 | -0.956 | 0.024 | 0.807 |
| MDA | -0.314 | -0.950 | 0.157 | 0.289 |
| **H<sub>2</sub>O<sub>2</sub>** | -0.505 | -0.863 | 0.610 | **0.003** |
| Nitrite | -0.307 | 0.952 | 0.054 | 0.647 |
| Nitrate | 0.414 | -0.910 | 0.029 | 0.767 |
| **Number of shoots** | 0.225 | 0.974 | 0.329 | **0.060** |
| Aboveground biomass | -0.385 | 0.923 | 0.110 | 0.450 |
| Belowground biomass | -0.981 | -0.192 | 0.062 | 0.639 |
| **Internode length** | -0.533 | -0.846 | 0.492 | **0.008** |

### 3.7 Core microbiome of native and invasive seagrass

We identified a total of 53 genera occurring in at least 80% of the samples that were shared among both native and invasive seagrass, representing their core microbiome (Figure 6A; Supplementary Table S5). Two genera were present in all groups, except when the invasive was growing alone, and were identified as an uncultured genus of the class *Alphaproteobacteria* and the genus *Aquibacter*. The native *Cymodocea nodosa* exhibited a core microbiome comprising 87 genera (Figure 6B; Supplementary Table S6), with no taxa unique to only one or two groups. In contrast, the invasive *H. stipulacea* harbored a smaller core of 24 genera (Figure 6C; Supplementary Table S7), and similarly, no taxa were exclusive to only one or two groups.

**Figure 6.**
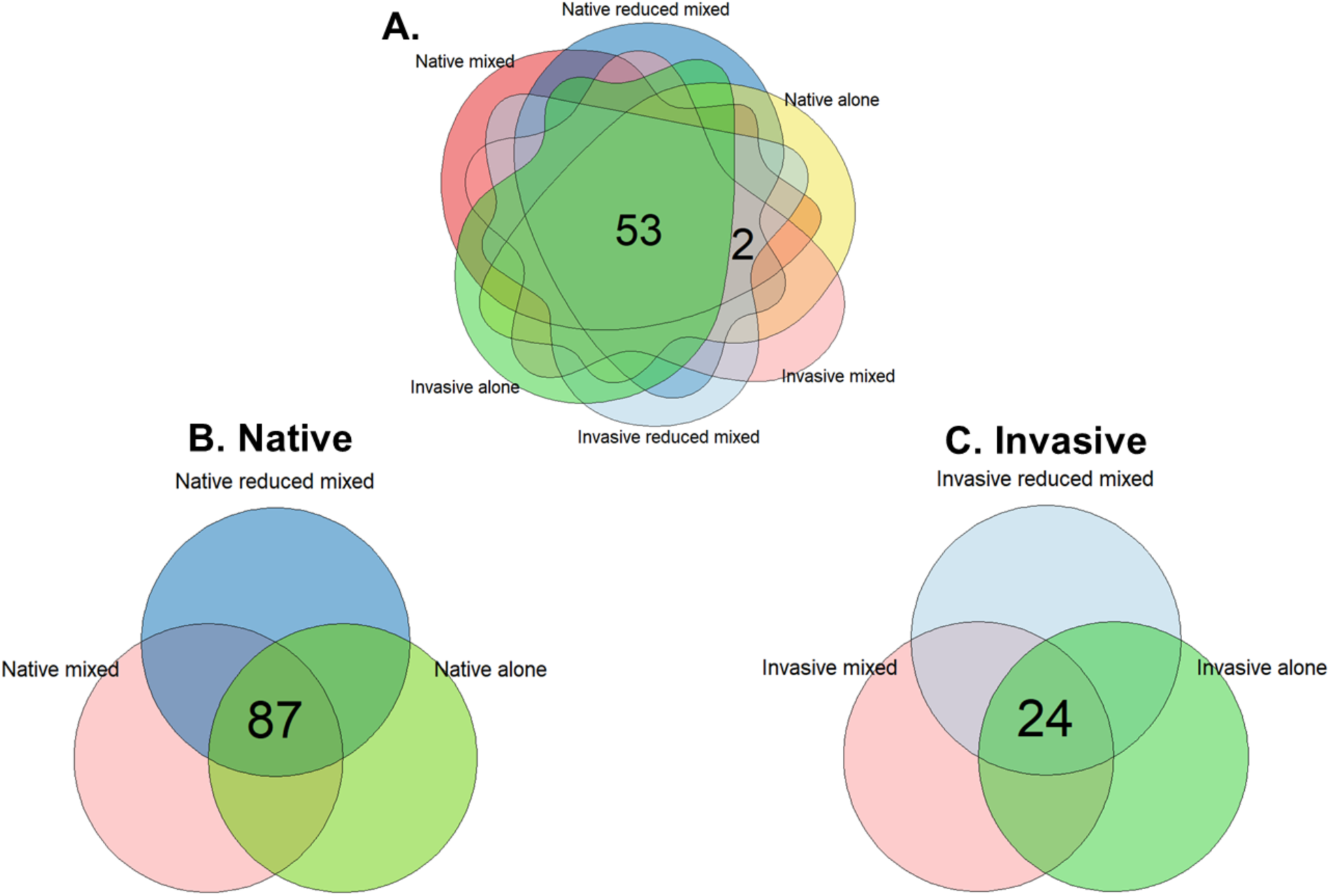
Venn diagrams of unique and concordant genera observed in the core microbiome of the seagrass samples. Empty cells indicate zero values. (A.) Unique and concordant genera observed in 80% of the combined native and invasive seagrass samples. (B.) Unique and concordant genera observed in 80% of the native *C. nodosa* samples only. (C.) Unique and concordant genera observed in 80% of the invasive *H. stipulacea* samples only.

Interestingly, we detected more differentially abundant bacterial genera of the core microbiome between native and invasive seagrass species when growing in mixed communities or alone (Figure 7A). Bacterial communities appeared more similar when the native seagrass grew in reduced mixed treatments and when it grew alone, than when grown with the invasive. This trend is the opposite for the invasive *H. stipulacea*, where bacterial communities looked similar to each other when grown in ambient and in reduced mixed treatments, compared to when grown alone.

**Figure 7.**
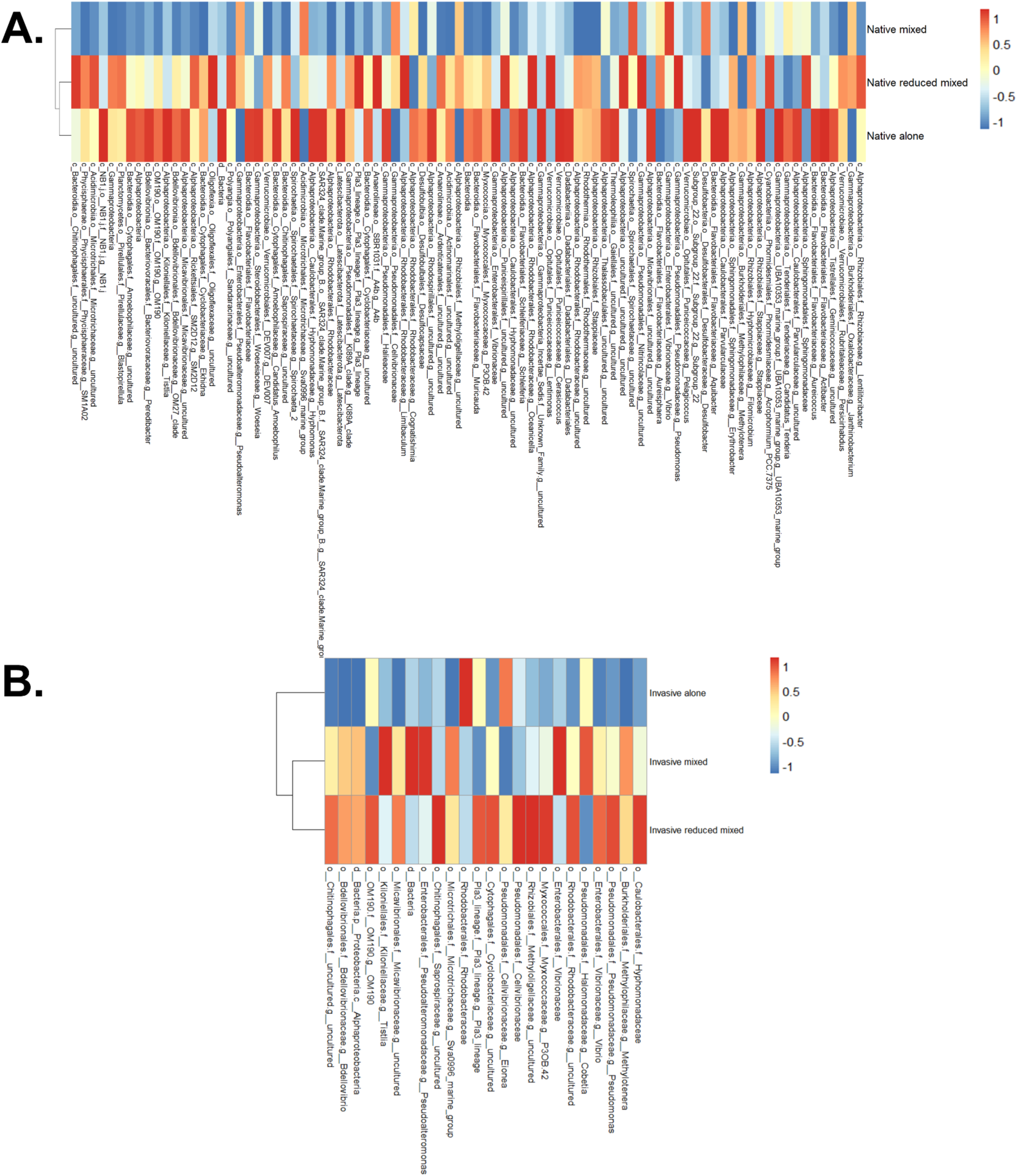
Heatmaps of genera observed in the core microbiome of native and invasive seagrass samples using Bray-Curtis distances and the Ward clustering algorithm on the genus level. A red color indicates a higher frequency of taxa (similarity), while a blue color indicates a lower frequency of taxa (dissimilarity). Hues indicate intensity. (A.) Genera observed in 80% of the combined native and invasive seagrass samples. (B.) Genera observed in 80% of the native *C. nodosa* samples only. (C.) Genera observed in 80% of the invasive *H. stipulacea* samples only.

In the core microbiome of the native seagrass *Cymodocea nodosa*, microbial activity resembled more closely when the native seagrass grew with the invasive *H. stipulacea* in reduced mixed densities and when it grew alone, compared to taxa when it grew in ambient conditions with the invasive *H. stipulacea* (Figure 7B). Over 35 taxa– such as *Bdellovibrio*, members of *Saprospiraceae*, *Cyclobacteriaceae*, and *Flavobacteriaceae–* were abundantly present when the native *C. nodosa* grows alone than when it grew mixed with the invasive in full or reduced densities.

In contrast, only a limited set of taxa—including *Pseudorhodobacter* and members of *Rhodobacteraceae*—showed higher abundance when growing at ambient densities with the invasive seagrass, suggesting that interspecific interactions may enhance the recruitment or activity of specific microbial associates. In the core microbiome of the invasive seagrass, many taxa that were highly abundant in mixed environments were absent when the invasive grew alone (Figure 7C). For example, *Bdellovibrio* and other members of the class *Alphaproteobacteria* showed increased activity in mixed conditions but were negatively associated when the invasive grew alone. Conversely, when the invasive species grew alone, we observed positive activity in *Rhodobacteraceae* and the genus *Eionea*, which were lower in abundance and grown in mixed and reduced mixed densities with the native seagrass.

## 4. Discussion

This is the first study to demonstrate that microbial communities shift significantly when mixed with *C. nodosa* and *H. stipulacea* Mediterranean beds than alone in monoculture, hinting at competition between native and invasive seagrass. Our results suggest that *H. stipulacea* may influence the outcomes of success, as *H. stipulacea* exhibited increased ASV richness in the presence of the native seagrass, while the native seagrass had a more diverse core microbiome and increased oxidative stress when grown alone, as well as lower ASV richness in the presence of the invasive seagrass. Importantly, combining these data with biochemical and morphological data from Chiquillo et al. (2022) suggests that microbes living on the phyllosphere positively influence the growth of invasive *H. stipulacea,* and likely contribute to its success as an invasive species.

Our results reveal an asymmetric pattern: *H. stipulacea*’s microbial diversity increased in the presence of *C. nodosa*, while *C. nodosa*’s microbial diversity decreased in the presence of *H. stipulacea*. Although much of the observed variation is primarily driven by species identity, rather than specific treatment contrasts, these patterns suggest that competition may directly reshape the structure and diversity of seagrass-associated microbes, with potential implications for invasion dynamics. Given that *H. stipulace*a thrives in both disturbed and open habitats and successfully invades intact native communities, we demonstrate that *H. stipulacea* negatively affects *C. nodosa* density and bacterial diversity, thus implying that antagonistic interactions with native seagrass may be a key aspect of *H. stipulacea’s* invasion ecology.

### 4.1 Native species, though diverse, may fail to resist invasion, at their own cost

We found that the native *C. nodosa* exhibited the highest microbial diversity in monoculture, along with a more diverse core microbiome. When grown with *H. stipulacea*, *C. nodosa*’s diversity declined, and plant density was reduced. Similar to Conte et al. (2023), we observed that *C. nodosa*’s highest diversity occurred when it grew alone, whereas in mixed assemblages, it exhibited reduced density, lower diversity, and higher phenol content, which is a biochemical indicator of stress (Mejia et al. 2016).

Our findings confirm that *C. nodosa’s* bacterial diversity is determined by its cellular stress and its desire to seek habitat away from the invasive *H. stipulacea* in mixed environments (e.g. internode length). The reasoning is that competition between plant species can reshape microbial communities, with consequences for native species fitness. Conte et al. (2023) also observed that *C. nodosa* boasted the highest bacterial diversity in monoculture, while *C. nodosa* reduced its density, depicted lower bacterial diversity, and had higher phenol content in syntopy with *H. stipulacea*. High phenol content is linked with stress in seagrasses and is equipped as a defense mechanism against threats such as predation, low light regimes, pollution, ocean acidification, and competition by marine macrophytes (Mejia et al. 2016). Conte et al.’s findings, alongside Chiquillo et al.’s (2022) observations that *C. nodosa* exhibited the greatest increase in shoot number when grown alone, suggest that *C. nodosa’s* associated microbial community may be ineffective at resisting invasion when co-occurring with the invasive seagrass. These shifts seem to mirror patterns seen in other systems where plant-plant competition reduces beneficial microbial associations (Foster & Bell 2012; Ke & Wan 2019), can lead to a loss of symbionts (Perreault & Laforest-Lapointe 2022), and stress physiology correlates with altered microbiomes, specifically where beneficial microbiomes related to plant tissues suppress stress (Zandi & Schnug 2022).

While the majority of studies focus on how microbes mediate competitive outcomes between plants, our approach examined how interspecific competition itself hosts differential microbial communities. Next steps would be to test for direct microbial transfer between species. Contrary to expectations, the native seagrass exhibited significantly higher oxidative stress (H₂O₂) levels compared to *H. stipulacea*. The capacity to scavenge reactive oxygen species such as H₂O₂ is a key determinant of a plant’s tolerance to environmental stress. Higher H₂O₂ has been observed in competitive superiors where elevated H₂O₂ concentrations were seen in monocultures compared to mixed assemblages (Ouyang et al. 2023; Navas et al. 1999); for example a noxious marine endophyte, *P. Crispus, which is* classified as a competitive superior, had higher H₂O₂ concentrations in monoculture than mixed (Chalanika et al. 2017). However, our results suggest that despite being competitively inferior to *H. stipulacea*, *C. nodosa’s* disproportionately high H₂O₂ levels may indicate that oxidative stress in this case reflects a competitive disadvantage rather than superiority.

### 4.2 Recruitment by the invasive species potentially drives its success

Competitive interactions between *H. stipulacea* and three other native seagrasses, *H. decipiens*, *S. filiforme,* and *T. testudinum* have shown displacement of natives from the Caribbean (Steiner & Willette 2015, Muthukrishnan et al 2020, Willette & Ambrose 2012), and its recent expansion of *H. stipulacea* into South Florida has been reported (Campbell et al. 2025). Not surprisingly, even when alone, our results support the theory that *H. stipulacea* is a driver of its own invasion success, possibly by actively shaping its microbial community and exploiting its environment to its advantage. In a “natural [field] experiment of seagrass interaction” in the Mediterranean, Conte et al. (2023) found evidence that *H. stipulacea’s* microbial communities are also plastic when growing with the native *C. nodosa*, displaying a high bacterial variability and supporting the host generalist hypothesis. In addition to negatively impacting the microbial community of native seagrass, results indicate that invasive *H. stipulacea* can drive its own success by recruiting beneficial bacteria in mixed environments (Supplementary Table S8) without the need for a disturbance to remove native seagrass species. It is possible that once *H. stipulacea* fragments dislodge from deeper meadows and encounter natives in shallower waters, higher nitrogen efficient use allows this species to rapidly colonize open or available space, even under oligotrophy. We know that *H. stipulacea* is a strong invader even in monoculture, where the species can colonize open habitat (Willette et al. 2014; Winters et al. 2020; Smulders et al. 2017). Like Conte et al., we found that *H. stipulacea* harbored a less diverse microbiome than *C. nodosa* in the Mediterranean Sea and that between the two species, *C. nodosa* hosted the richest and most abundant microbiome in monoculture, while *H. stipulacea* bacterial diversity was highest in mixed environments. *Rhodobacteraceae* sp. might explain the ability of *H. stipulacea* to expand into uncolonized areas and shallow water habitats, as they are capable of assisting with carbon and sulfur cycling as well as photosynthesis in marine anaerobic environments (Pujalte et al. 2014; Crump et al. 2018), producing antibacterial compounds (Hurtado-McCormick et al. 2019), and are potentially involved in biofilm formation (Vogel et al. 2020; Conte et al. 2023). They have been frequently identified in seagrasses, including *H. stipulacea* (Mejia et al. 2016; Rotini et al. 2017; Stuij 2018; Hurtado-McCormick et al. 2019; Garcias-Bonet et al. 2021; Ling et al. 2021; Conte et al. 2023) and have been associated with *H. stipulacea* leaves in high light and hydrodynamic regimes (Mejia et al. 2016).

Results from controlled mesocosms in the Mediterranean Sea add to the existing knowledge of *H. stipulacea* microbial diversity. We observed lower microbial community diversity compared to previous observations in both mixed and monoculture environments across the Red Sea (Wahbeh et al. 1984; Pereg et al. 1994; Weidner et al. 1996, 2000; Mejia et al. 2016; Rotini et al. 2017; Garcias-Bonet et al. 2020; Abdel-Wahab et al. 2021), Mediterranean (Conte et al. 2021, 2023), and Caribbean Seas (Stuij 2018; Aires et al. 2021). This contrast may be due to the experimental context of our study, as samples were collected from a mesocosm six weeks after transplantation, rather than directly from natural donor sites. Although we did not detect the same taxa, our findings align with Conte et al. (2023), who identified beneficial *H. stipulacea* bacteria in the Mediterranean Sea such as the nitrogen-fixing *Microcystaceae*, *Nisaeaceae*, *Phormidiaceae*; sulfur-fixing *Desulfocapsacaceae*; and leaf colonizing *Pirellulaceae*. Our results similarly align with Aires et al. (2021), who identified beneficial *H. stipulacea* bacteria in the Caribbean Sea such as the plant growth-promoting *Halomonas*, quorum-inhibiting *Salinimonas* and *Salinicola*, heavy metal-tolerant *Lysinibacillus*, and halotolerant genera such as *Rhodococcus* and *Azospirillum*, which support seed germination under high salinity.

Due to constraints, our study examined only phyllosphere bacteria associated with the native and invasive seagrass, although it is widely accepted that the rhizosphere exhibits the richer microbial community (Rotini et al. 2017; Li et al. 2025). It is likely that rhizosphere bacteria also play important roles in *H. stipulacea* invasion success, particularly since seagrass above- and belowground compartments possess distinct microbes (Stuij 2018; Winters et al. 2020; Aires et al. 2021) and sediment microbes can greatly impact nutrient acquisition and other critical plant processes.

### 4.3. Alternative hypotheses: Microbes as drivers or passengers of competitive success

One possible explanation for reduced microbial diversity in *C. nodosa* when grown with *H. stipulacea* is allelopathy. *H. stipulacea* could harbor microbial or fungal associates (Gribben et al. 2017) that produce compounds negatively affecting the native species (Orr et al. 2005). For example, we found a strong positive association between *H. stipulacea* in intraspecific competition and the genus *Perspicuibacter*, involved in sulfur cycling, in contrast to mixed assemblages. This pattern differs from studies reporting strong negative effects of interspecific competition among seagrasses. We also observed a positive association between *H. stipulacea* grown alone and members of *Rhodobacteraceae*. While allelopathy is well documented in terrestrial plants (Bais et al. 2003; Orr et al. 2005; Chengxu et al. 2011; Greer et al. 2015; Chen et al. 2017; Kalisz et al. 2020) and in some seagrass–macroalgae interactions (Dumay et al. 2004; Pergent et al. 2008; Raniello et al. 2007), to our knowledge, seagrass–seagrass allelopathy has not been reported. *Vibrio*, significantly more associated with *C. nodosa* in mixed assemblages, is notable for its algicidal (Sakata et al. 2011; Setiabundi et al. 2023), antimicrobial (Millan 2022; Wietz et al. 2010; Weitz et al. 2013; Romero-González et al. 2023; Alattas et al. 2024), and pathogenic properties (Sampaio et al. 2022; Alattas et al. 2024), as well as its production of siderophores—iron-chelating compounds with potential allelopathic effects (Wietz et al. 2013; Alattas et al. 2024; Dar et al. 2024).

Another possible mechanism is that *H. stipulacea* disrupts the phyllosphere community of *C. nodosa* in mixed assemblages. *Desulfobulbus*, the most significantly associated genus with *C. nodosa* grown alone (B = 1.107, q = 0.00214), is a sulfate-reducer common in seagrass holobionts and sediments (Küsel et al. 1999; Ling et al. 2021; Zhou et al. 2021). It has been linked to nutrient cycling, hydrocarbon degradation, and nitrogen fixation (Zhou et al. 2021; Wang et al. 2025). Other taxa significantly associated with *C. nodosa* alone include *Bermanella*, known for hydrocarbon degradation (Hu et al. 2017; Ribicic et al. 2018; Peeb et al. 2022; Howe et al. 2024), and the PS1 clade, related to nitrogen-fixing *Rhizobiales* adapted to low-nutrient conditions (Jimenez-Infante et al. 2014; Stuij 2018; Zhai et al. 2022). Loss of such taxa in mixed assemblages could reduce the biochemical processes that support native seagrass performance.

For *H. stipulacea*, several taxa were less associated when grown alone than in mixed assemblages, suggesting a capacity to retain beneficial microbes under different competitive scenarios. These included *Zeaxanthinibacter* (photoprotective pigment production), *Tenacibaculum* (organic matter degradation), *Photobacterium* (nitrogen fixation, nutrient provisioning), *Rhodospirillales* (anaerobic photosynthesis), and *Micavibrionales* (potential oil-degrader interactions). While 16S rRNA sequencing does not confirm functional activity, the taxonomic affiliations are consistent with traits that could enhance *H. stipulacea* growth and resilience. We also did not account for archaea, protists, viruses or fungal communities, which are known to mediate nutrient acquisition, host–microbe interactions, and organismal expansion (Van der Putten et al. 2007). As climate change is expected to promote the expansion of *H. stipulacea* in the Mediterranean (Winters et al. 2020; Beca-Carretero et al. 2023; Nguyen et al. 2020), understanding whether these microbial shifts are a cause or a consequence of competitive success remains a critical next step.

### 4.4 Concluding statement

This study illuminates how microbes might facilitate the growth and invasion success of the invasive seagrass *Halophila stipulacea* in mixed communities with the native *Cymodocea nodosa* in the Mediterranean Sea. Specifically, we show how microbial assemblies shift with competition. When competition alters invasive performance, microbial communities also differ. Ultimately, our results support the association between *H. stipulacea* phyllosphere bacteria and its invasion success in mixed Mediterranean environments; however, further research is necessary into the mechanisms behind this association. Specifically, investigations into whether *H. stipulacea* selectively recruits beneficial microbes, facilitates its own invasion via microbe-mediated allelopathic interactions, or is passively shaped by microbial community assembly processes is required. Overall, we present valuable new insights into plant-microbe interactions in marine ecosystems and the possible benefits of seagrass-microbe interactions in *H. stipulacea* invasions.

## Author Contributions

Conceptualization KLC PHB; Methodology KLC, MIV, VF, CA KD; Validation SMC, VF; Formal Analysis: SMC, EHF, FGV; Investigation KLC, MIV; Resources VF, PHB, MIV, FGV; Data Curation EHF, KLC, FGV, SMC; Software: SMC, EHF; Writing—Original Draft Preparation EHF, KLC; Writing—Review & Editing EHF, PHB, MIV, KLC, SMC; Validation: SMC, Visualization EHF; Supervision PHB and KLC; Project Administration KLC; Funding Acquisition PHB, KLC, FGV. All authors have read and agreed to the published version of the manuscript.

## Data Availability Statement

The raw 16s rRNA sequences of this study are openly available on NCBI- Accession: PRJNA1291277. Data can be accessible via OSF, and the link for peer review: https://osf.io/zg8xm/overview. All data will be made accessible to public when published.

## Acknowledgements

We thank Stylianos Kanakaris for his support in field and lab work. We would like to acknowledge Juanse Ramirez and Jose Enrique Arrarias for their immense support in funding and providing feedback.

## Supplemental Materials

**S1. Supplementary Table 1:** Linear Mixed Effects model of Chlorophyll a, Chlorophyll b, Carotenoids, MDA and H_2_O_2_. Numerator (numDF) and denominator (denDF), F-value and p-value are summarized.

| **Variable** | **Parameter** | **numdf** | **dendf** | **F-value** | **Pr(>F)** |
| --- | --- | --- | --- | --- | --- |
| **Chl-a** | Intercept | 1 | 43 | 771.68 | **<.0001** |
|  | Treatment | 2 | 43 | 2.8394 | 0.0695 |
|  | Species | 1 | 43 | 44.0615 | **<.0001** |
|  | Treatment x<br>Species | 2 | 43 | 6.2257 | **0.0042** |
| **Chl-b** | Intercept | 1 | 43 | 600.94 | **<.0001** |
|  | Treatment | 2 | 43 | 1.0949 | 0.3437 |
|  | Species | 1 | 43 | 11.3188 | **0.0016** |
|  | Treatment x<br>Species | 2 | 43 | 4.4499 | **0.0175** |
| **Carotenoids** | Intercept | 1 | 43 | 716.147 | **<.0001** |
|  | Treatment | 2 | 43 | 12.949 | **<.0001** |
|  | Species | 1 | 43 | 3.5764 | 0.0654 |
|  | Treatment x<br>Species | 2 | 43 | 6.6622 | **0.003** |
| **H<sub>2</sub>O<sub>2</sub>** | Intercept | 1 | 12 | 282.635 | **<.0001** |
|  | Treatment | 2 | 12 | 3.03215 | 0.0859 |
|  | Species | 1 | 12 | 155.397 | **<.0001** |
|  | Treatment x<br>Species | 2 | 12 | 1.37104 | 0.2909 |
| **MDA** | Intercept | 1 | 12 | 956.557 | **<.0001** |
|  | Treatment | 2 | 12 | 1.5106 | 0.2599 |
|  | Species | 1 | 12 | 6.0007 | **0.0306** |
|  | Treatment x<br>Species | 2 | 12 | 3.8989 | **0.0496** |
| **Internode**<br>**length** | Intercept | 1 | 43 | 97.94588 | **<.0001** |
|  | Treatment | 2 | 43 | 1.67859 | 0.7697 |
|  | Species | 1 | 43 | 13.07934 | **<.0001** |
|  | Treatment x<br>Species | 2 | 43 | 1.15488 | 0.5478 |
| **NO2** | Intercept | 1 | 28 | 1509.2 | **<.0001** |
|  | Treatment | 3 | 28 | 1.4424 | 0.2515 |
| **NO3** | Intercept | 1 | 28 | 1509.2 | **<.0001** |
|  | Treatment | 3 | 28 | 1.4424 | 0.2515 |

**S2. Supplementary Table 2.** Kruskal-Wallis statistics for Shannon alpha diversity. Significant values are bolded.

| Kruskal-Wallis Test | X<sup>2</sup> | df | P |
| --- | --- | --- | --- |
|  | 7.745 | 5 | 0.171 |

**S3. Supplementary Table 3.**
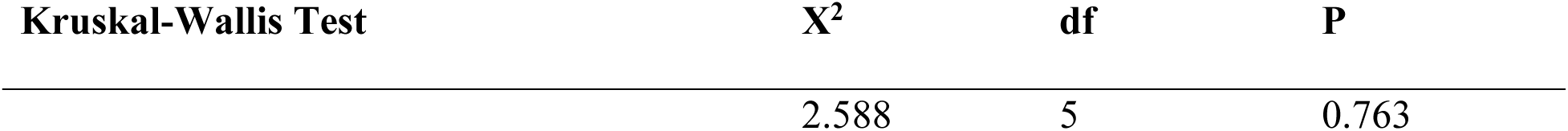
Kruskal-Wallis statistics for Pielou evenness (alpha diversity).

| Kruskal-Wallis Test | X<sup>2</sup> | df | P |
| --- | --- | --- | --- |
|  | 2.588 | 5 | 0.763 |

**S4. Supplementary Table 4.**
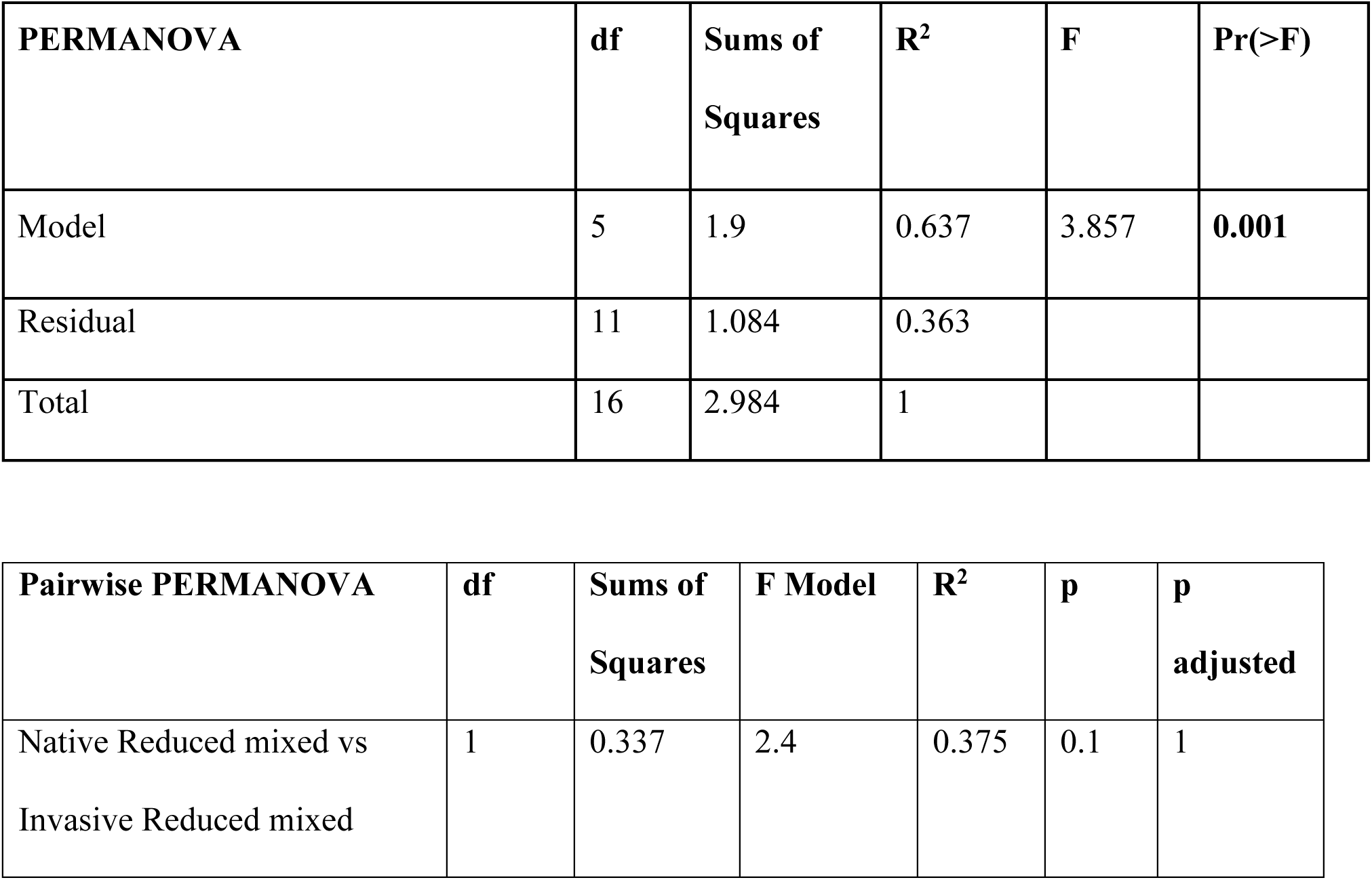
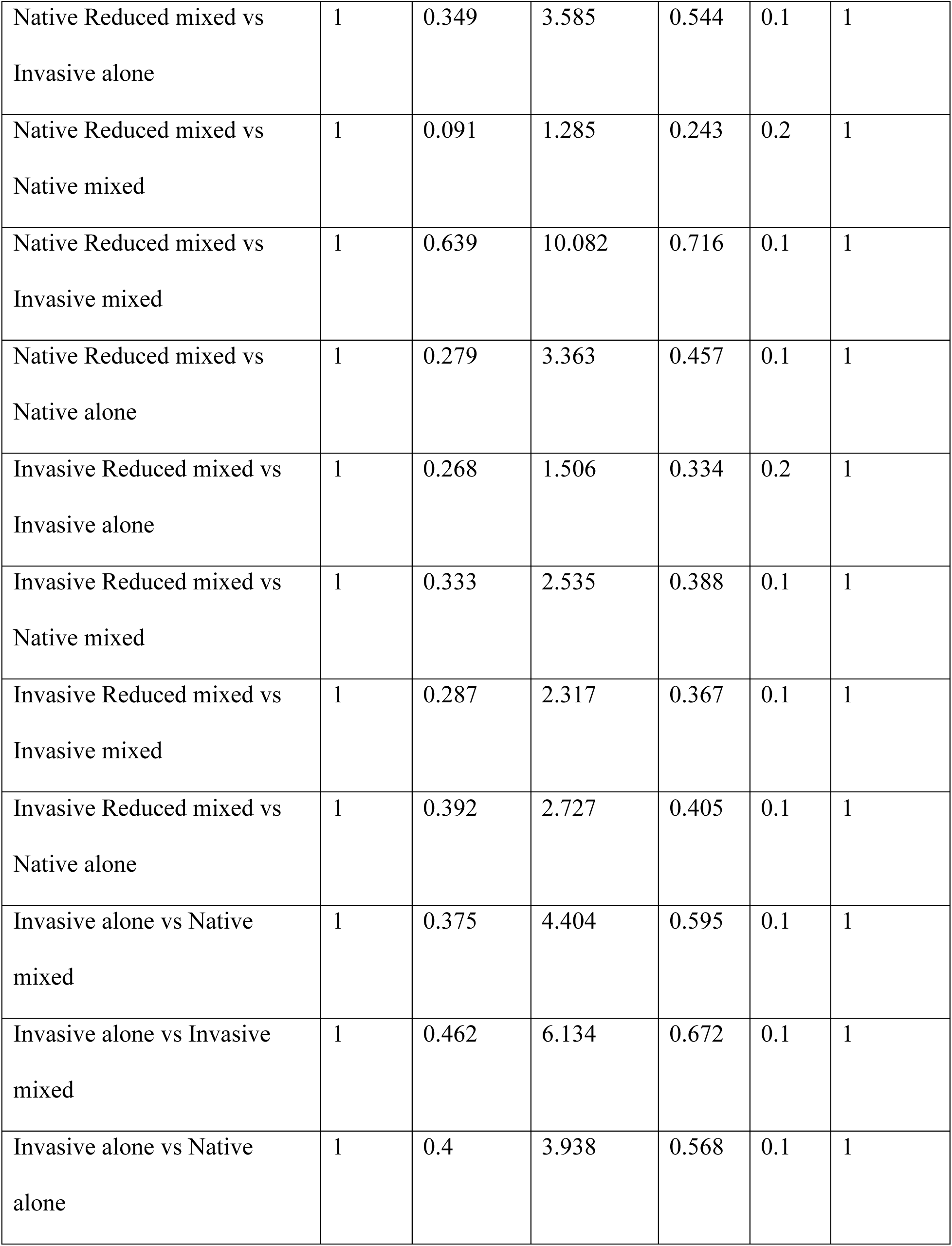
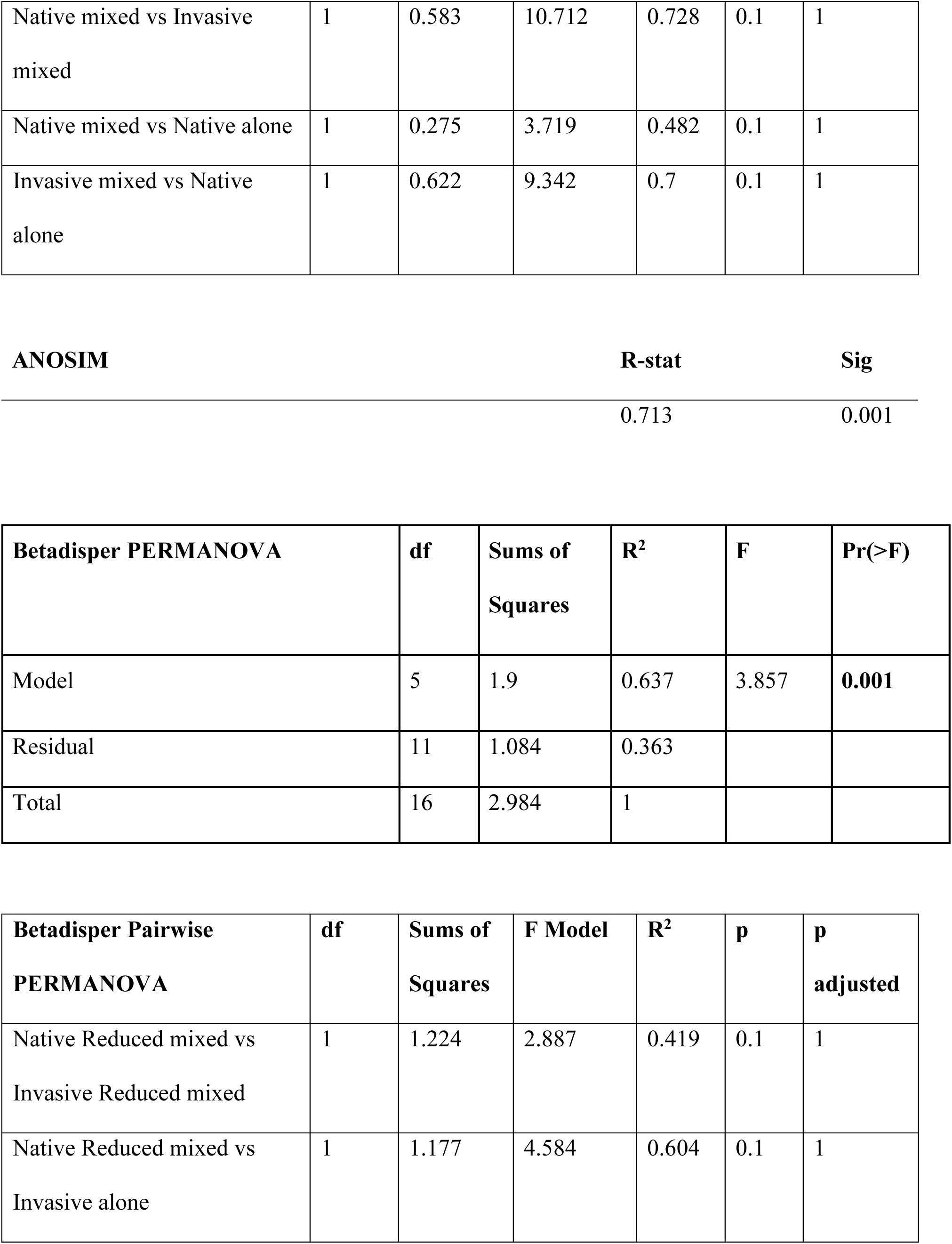
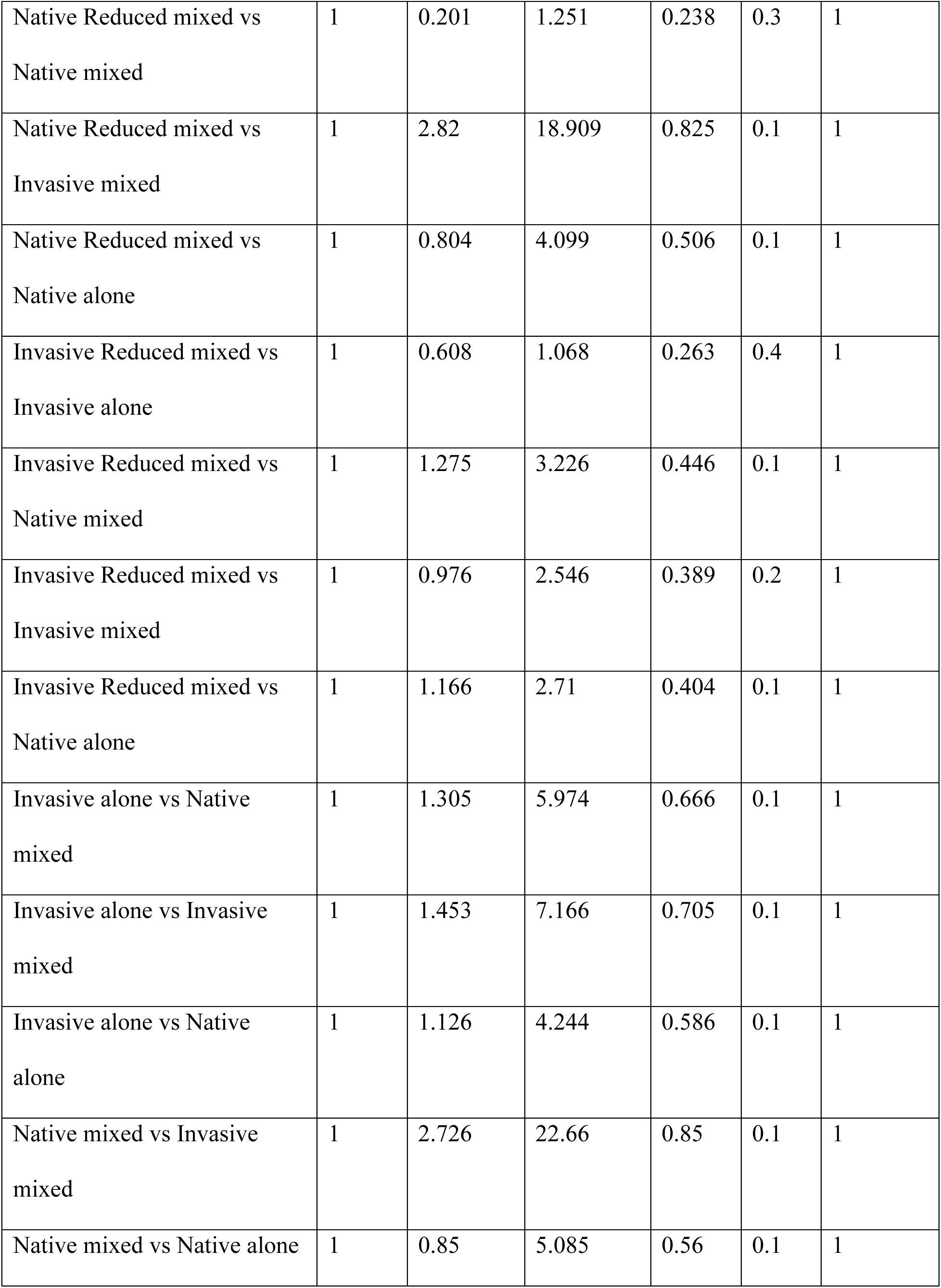
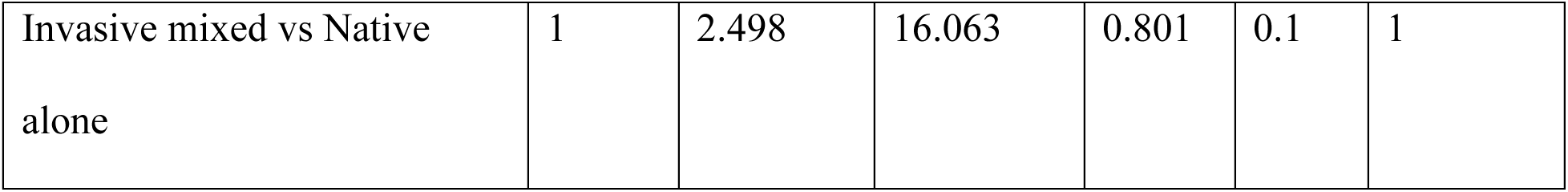
PERMANOVA, pairwise PERMANOVA, ANOSIM, and betadisper statistics for beta diversity. Significant values are bolded.

| PERMANOVA | df | Sums of Squares | R<sup>2</sup> | F | Pr(>F) |
| --- | --- | --- | --- | --- | --- |
| Model | 5 | 1.9 | 0.637 | 3.857 | **0.001** |
| Residual | 11 | 1.084 | 0.363 |  |  |
| Total | 16 | 2.984 | 1 |  |  |

| Pairwise PERMANOVA | df | Sums of Squares | F Model | R<sup>2</sup> | p | p adjusted |
| --- | --- | --- | --- | --- | --- | --- |
| Native Reduced mixed vs<br>Invasive Reduced mixed | 1 | 0.337 | 2.4 | 0.375 | 0.1 | 1 |
| Native Reduced mixed vs<br>Invasive alone | 1 | 0.349 | 3.585 | 0.544 | 0.1 | 1 |
| Native Reduced mixed vs<br>Native mixed | 1 | 0.091 | 1.285 | 0.243 | 0.2 | 1 |
| Native Reduced mixed vs<br>Invasive mixed | 1 | 0.639 | 10.082 | 0.716 | 0.1 | 1 |
| Native Reduced mixed vs<br>Native alone | 1 | 0.279 | 3.363 | 0.457 | 0.1 | 1 |
| Invasive Reduced mixed vs<br>Invasive alone | 1 | 0.268 | 1.506 | 0.334 | 0.2 | 1 |
| Invasive Reduced mixed vs<br>Native mixed | 1 | 0.333 | 2.535 | 0.388 | 0.1 | 1 |
| Invasive Reduced mixed vs<br>Invasive mixed | 1 | 0.287 | 2.317 | 0.367 | 0.1 | 1 |
| Invasive Reduced mixed vs<br>Native alone | 1 | 0.392 | 2.727 | 0.405 | 0.1 | 1 |
| Invasive alone vs Native<br>mixed | 1 | 0.375 | 4.404 | 0.595 | 0.1 | 1 |
| Invasive alone vs Invasive<br>mixed | 1 | 0.462 | 6.134 | 0.672 | 0.1 | 1 |
| Invasive alone vs Native<br>alone | 1 | 0.4 | 3.938 | 0.568 | 0.1 | 1 |
| Native mixed vs Invasive mixed | 1 | 0.583 | 10.712 | 0.728 | 0.1 | 1 |
| Native mixed vs Native alone | 1 | 0.275 | 3.719 | 0.482 | 0.1 | 1 |
| Invasive mixed vs Native alone | 1 | 0.622 | 9.342 | 0.7 | 0.1 | 1 |

|  |  |  |
| --- | --- | --- |
| **ANOSIM** | **R-stat** | **Sig** |
|  | 0.713 | 0.001 |
| **Betadisper PERMANOVA** | **df** | **Sums of Squares** | **R<sup>2</sup>** | **F** | **Pr(>F)** |
| Model | 5 | 1.9 | 0.637 | 3.857 | **0.001** |
| Residual | 11 | 1.084 | 0.363 |  |  |
| Total | 16 | 2.984 | 1 |  |  |

|  |  |  |  |  |  |  |
| --- | --- | --- | --- | --- | --- | --- |
| **Betadisper Pairwise PERMANOVA** | **df** | **Sums of Squares** | **F Model** | **R<sup>2</sup>** | **p** | **p adjusted** |
| Native Reduced mixed vs Invasive Reduced mixed | 1 | 1.224 | 2.887 | 0.419 | 0.1 | 1 |
| Native Reduced mixed vs Invasive alone | 1 | 1.177 | 4.584 | 0.604 | 0.1 | 1 |
| Native Reduced mixed vs<br>Native mixed | 1 | 0.201 | 1.251 | 0.238 | 0.3 | 1 |
| Native Reduced mixed vs<br>Invasive mixed | 1 | 2.82 | 18.909 | 0.825 | 0.1 | 1 |
| Native Reduced mixed vs<br>Native alone | 1 | 0.804 | 4.099 | 0.506 | 0.1 | 1 |
| Invasive Reduced mixed vs<br>Invasive alone | 1 | 0.608 | 1.068 | 0.263 | 0.4 | 1 |
| Invasive Reduced mixed vs<br>Native mixed | 1 | 1.275 | 3.226 | 0.446 | 0.1 | 1 |
| Invasive Reduced mixed vs<br>Invasive mixed | 1 | 0.976 | 2.546 | 0.389 | 0.2 | 1 |
| Invasive Reduced mixed vs<br>Native alone | 1 | 1.166 | 2.71 | 0.404 | 0.1 | 1 |
| Invasive alone vs Native<br>mixed | 1 | 1.305 | 5.974 | 0.666 | 0.1 | 1 |
| Invasive alone vs Invasive<br>mixed | 1 | 1.453 | 7.166 | 0.705 | 0.1 | 1 |
| Invasive alone vs Native<br>alone | 1 | 1.126 | 4.244 | 0.586 | 0.1 | 1 |
| Native mixed vs Invasive<br>mixed | 1 | 2.726 | 22.66 | 0.85 | 0.1 | 1 |
| Native mixed vs Native alone | 1 | 0.85 | 5.085 | 0.56 | 0.1 | 1 |
| Invasive mixed vs Native<br>alone | 1 | 2.498 | 16.063 | 0.801 | 0.1 | 1 |

**S5. Supplementary Table 5.**
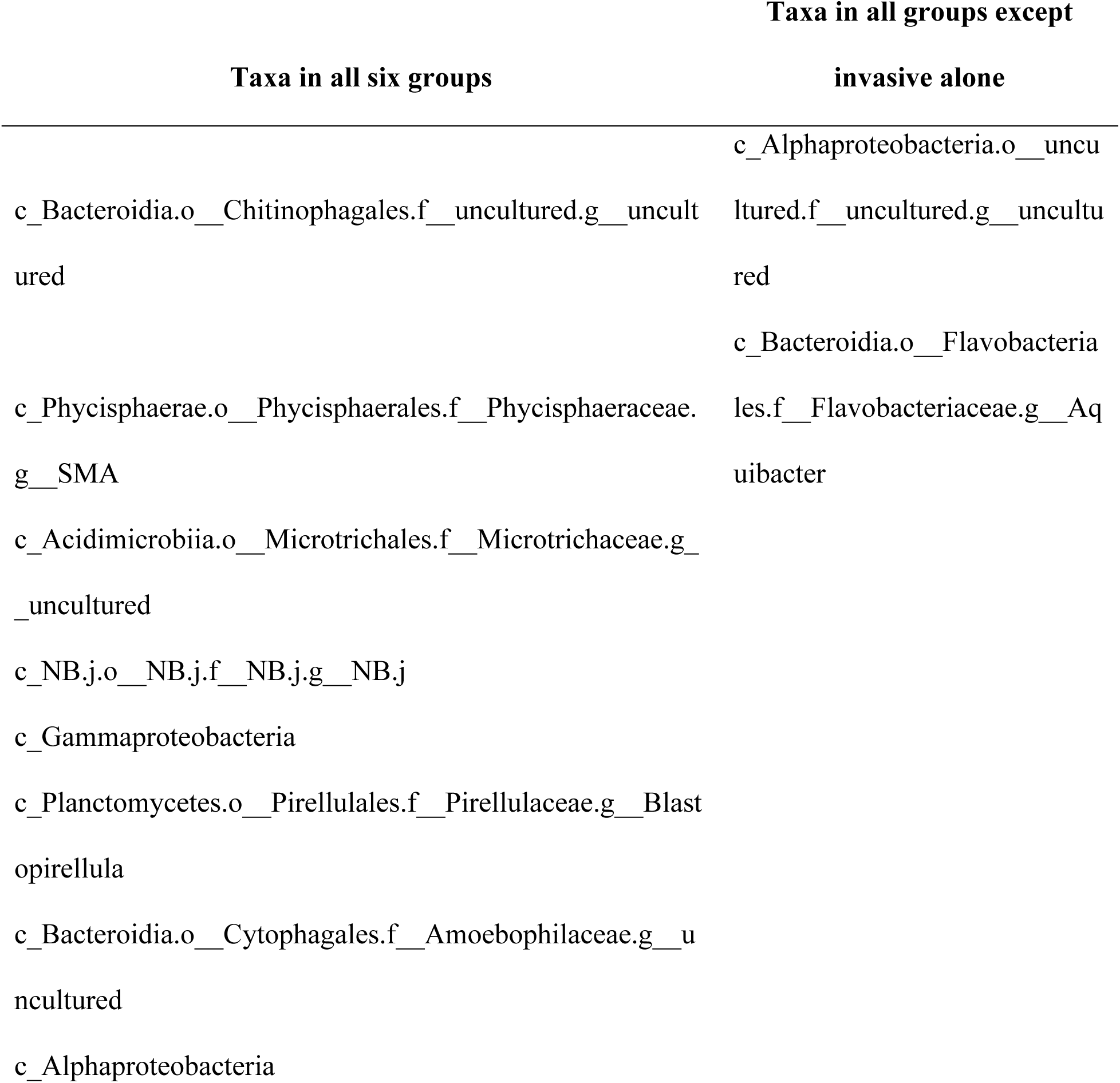
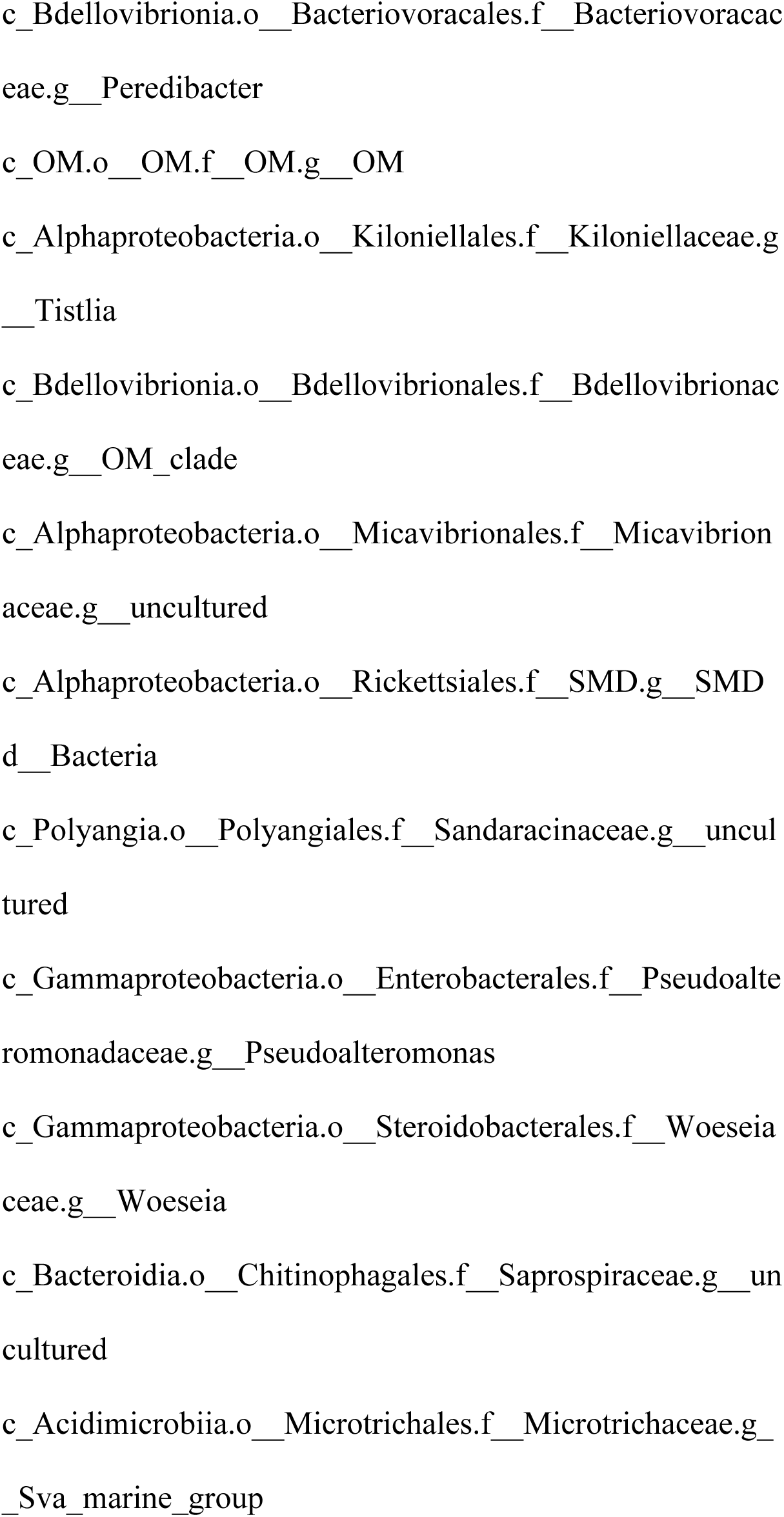
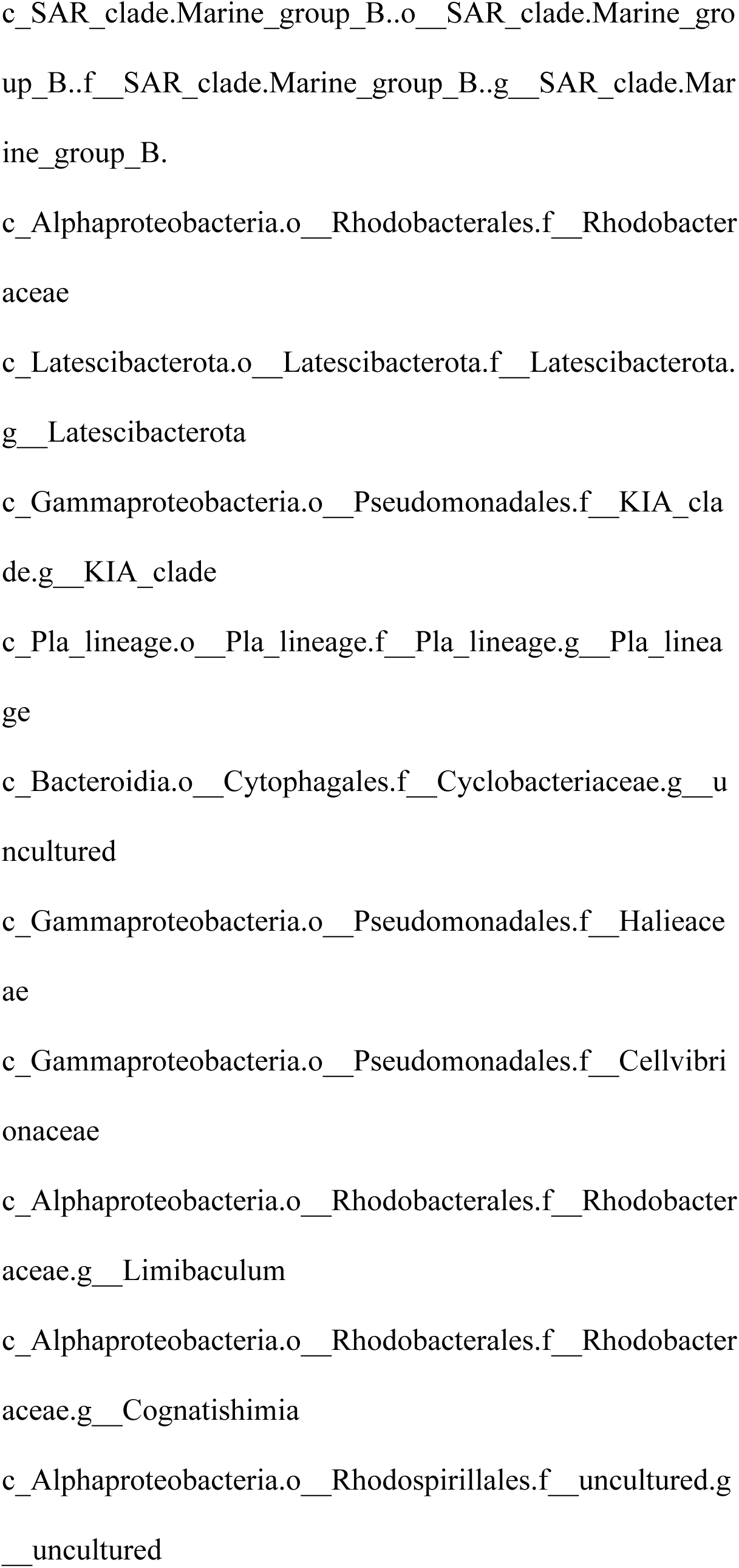
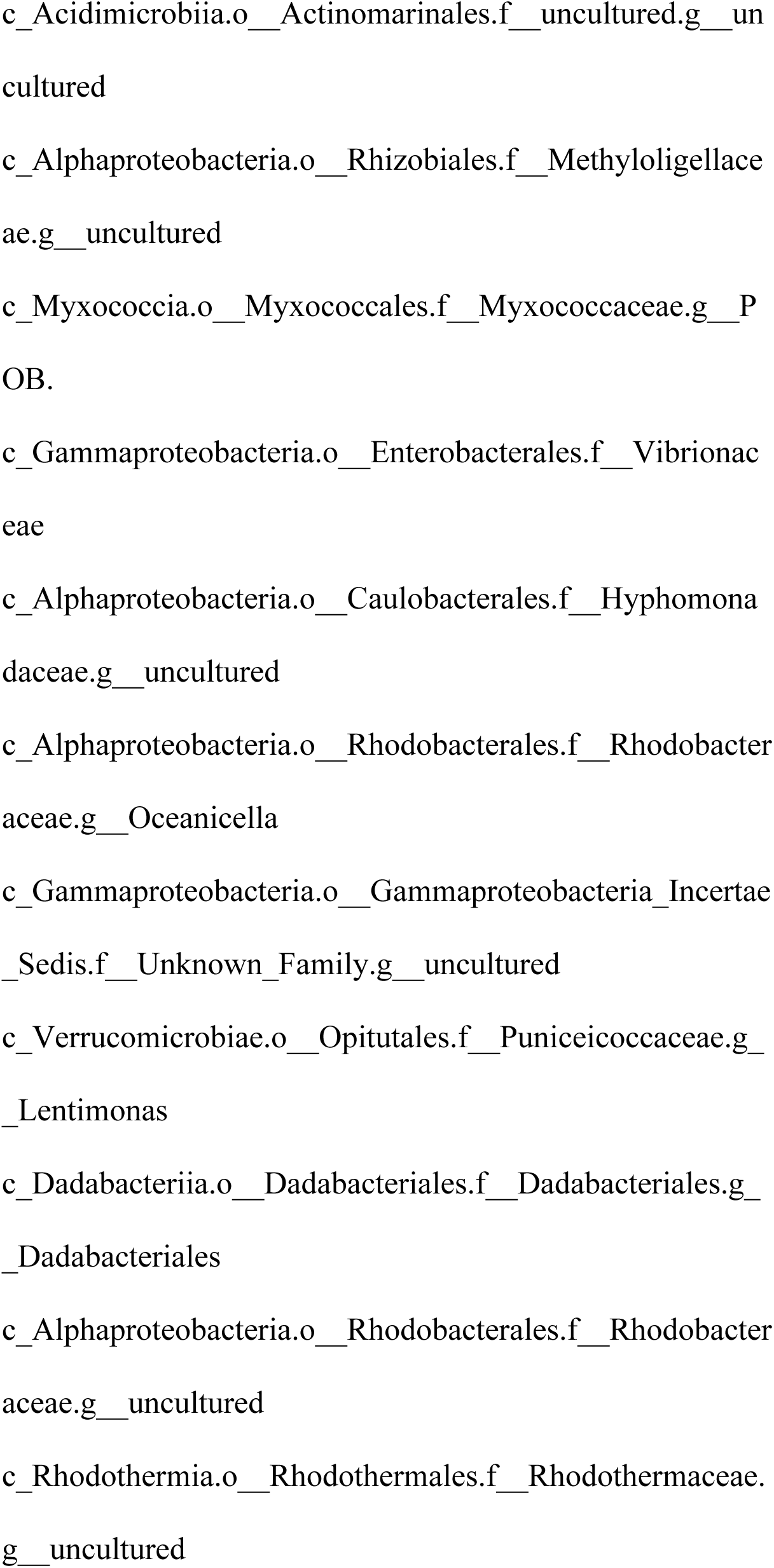
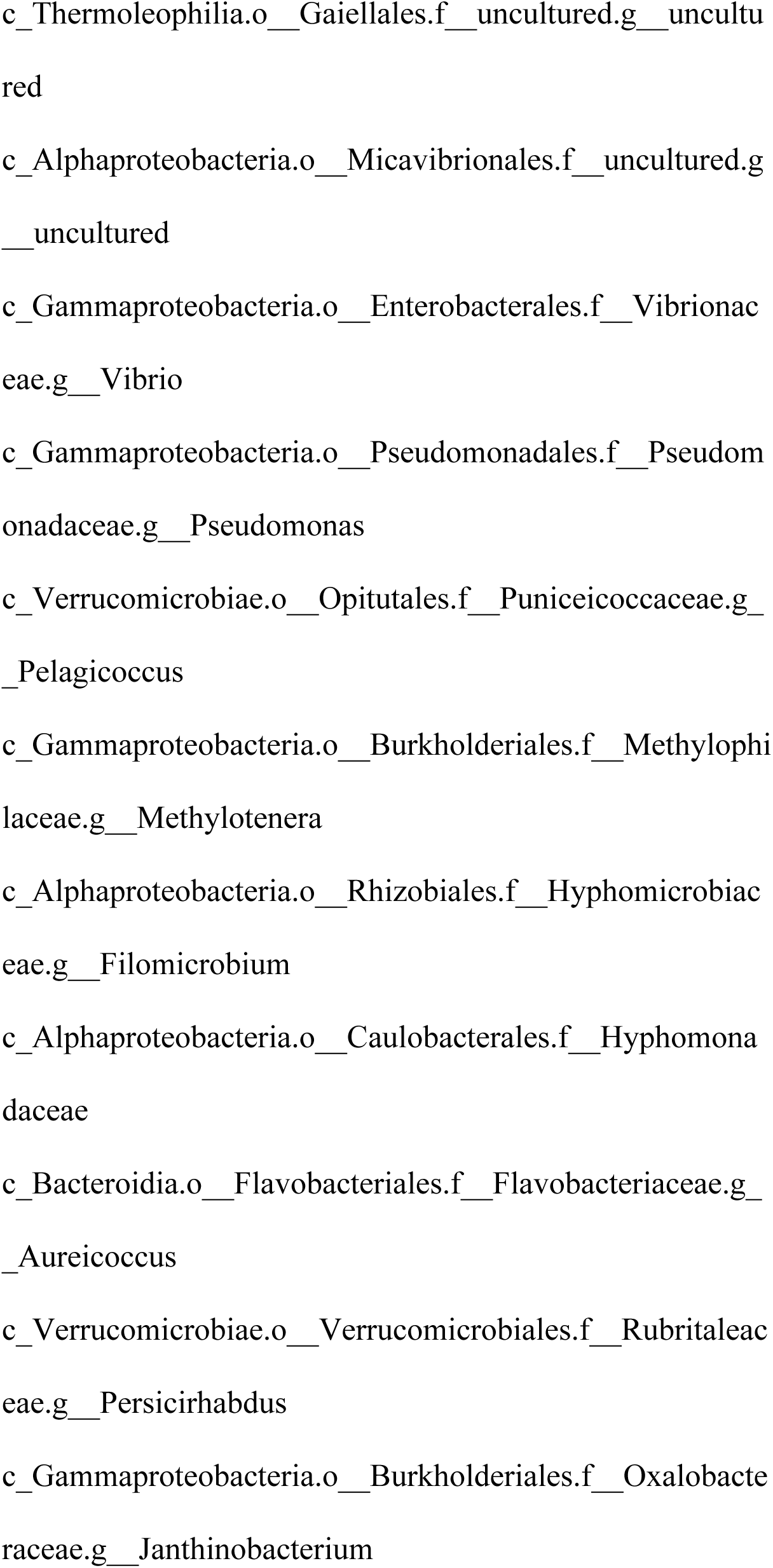
Table of core taxa shared among all six groups and all groups except invasive alone (see Figure 6A).

**S6. Supplementary Table 6.**
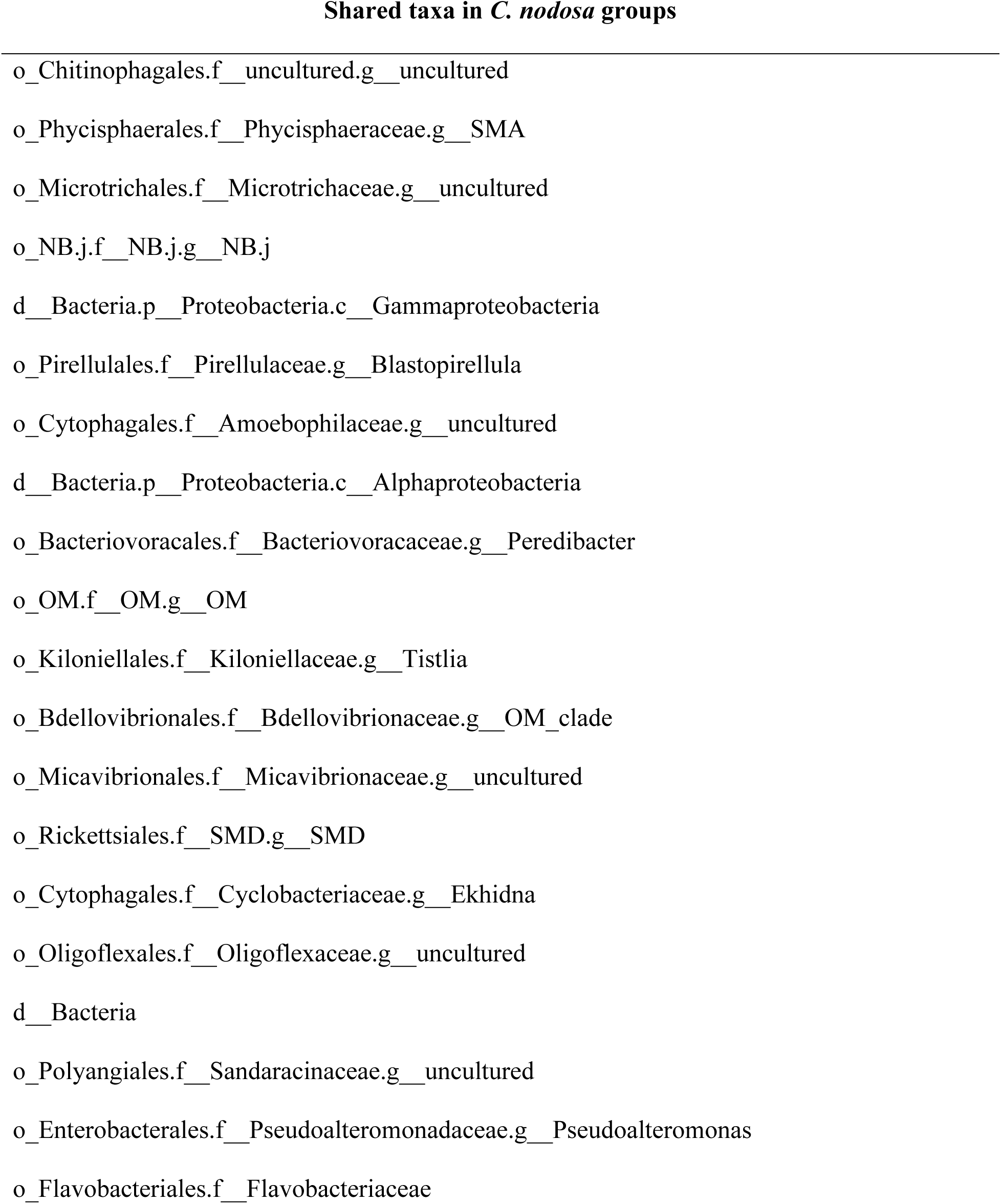
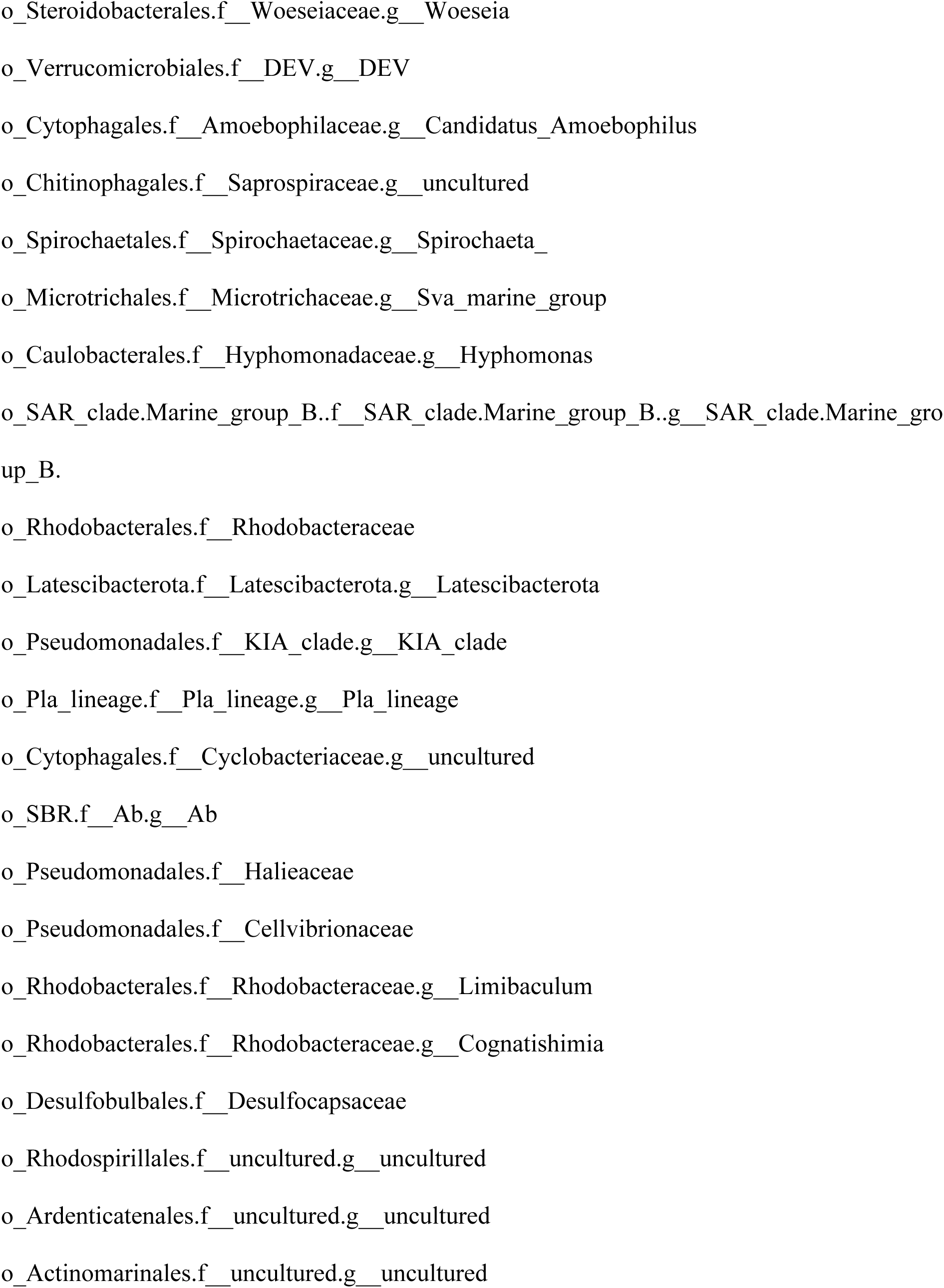
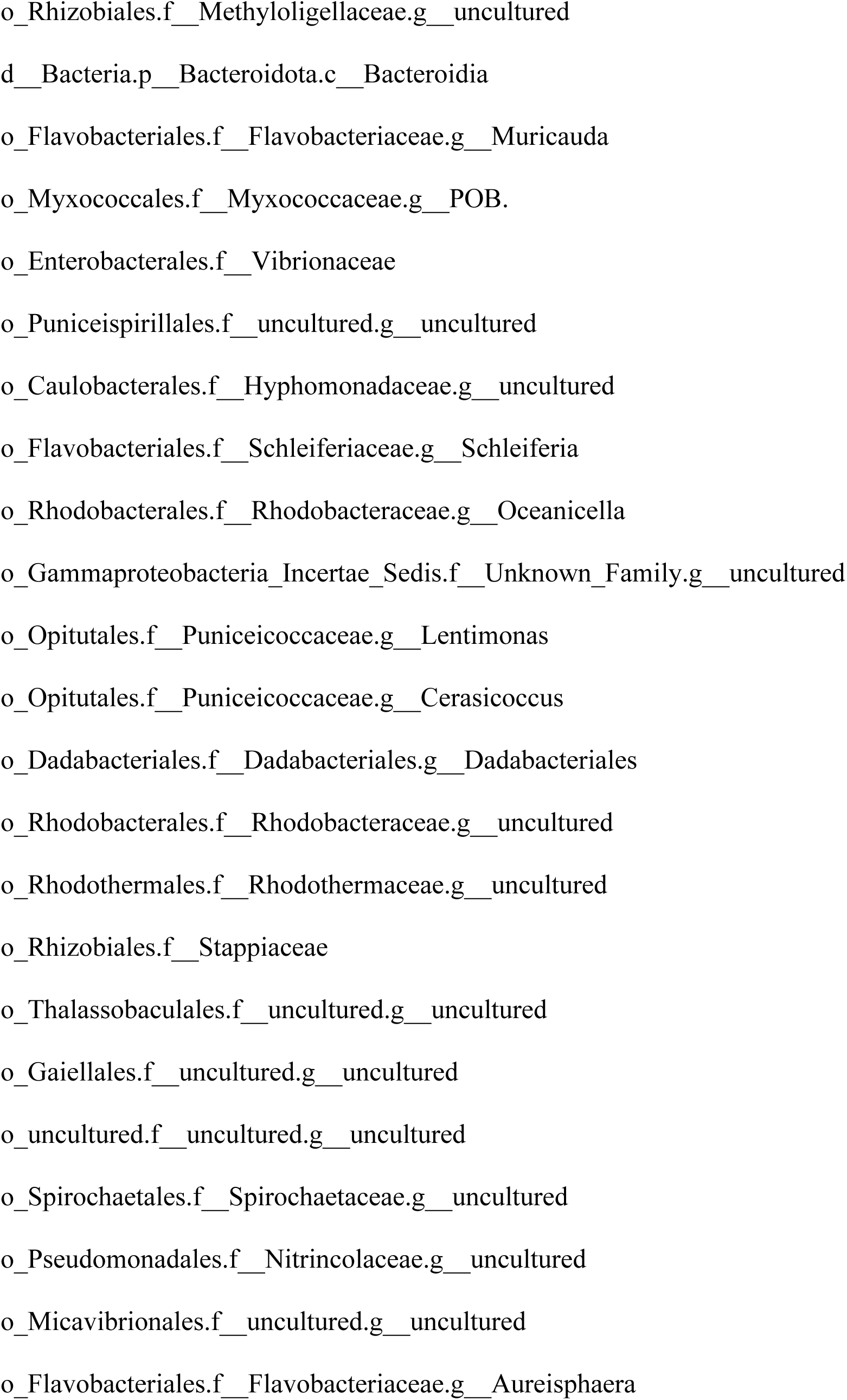
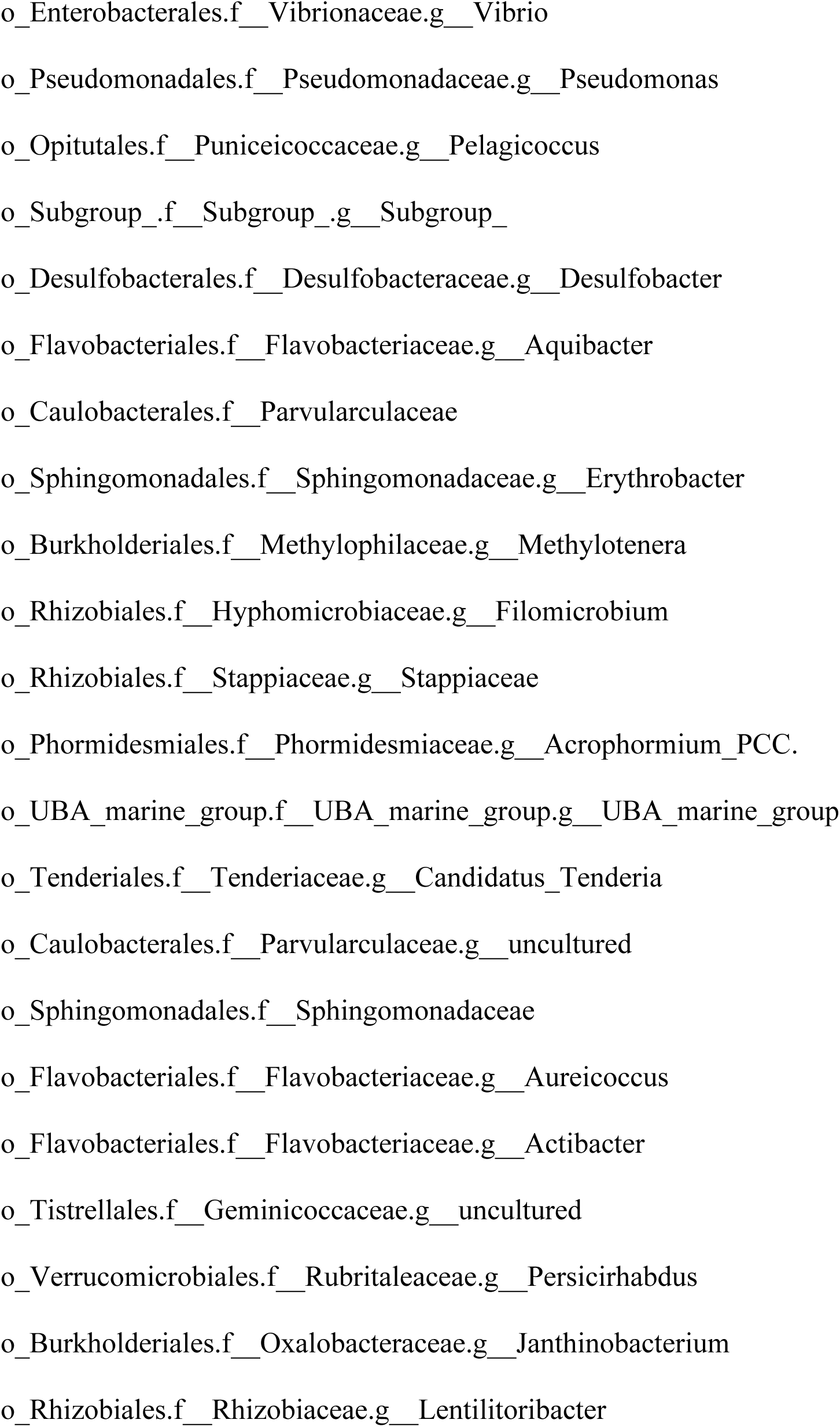
Table of core taxa shared among *C. nodosa* groups (see Figure 6B).

**S7. Supplementary Table 7.**
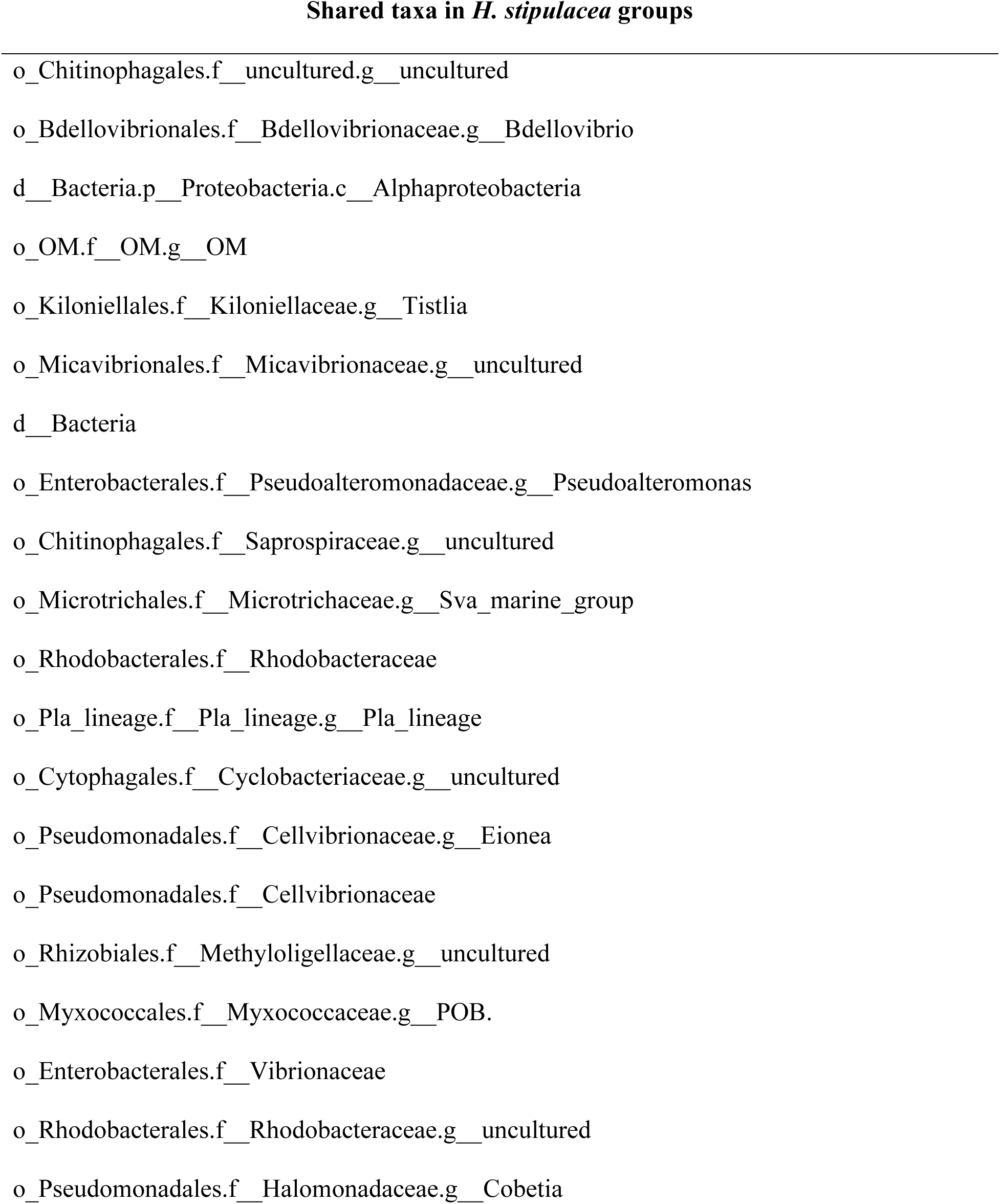
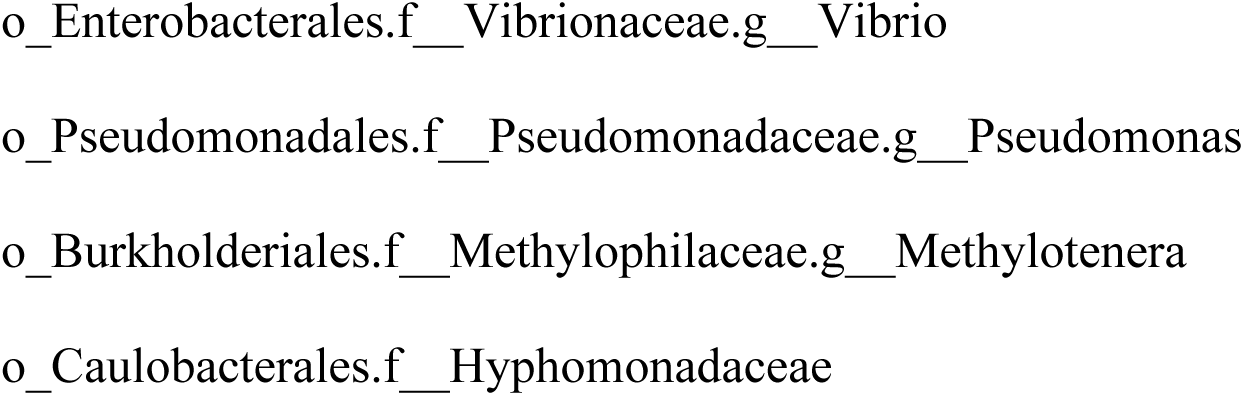
Table of core taxa shared among *H. stipulacea* treatments (see Figure 6C).

**S8. Supplementary Table 8.**
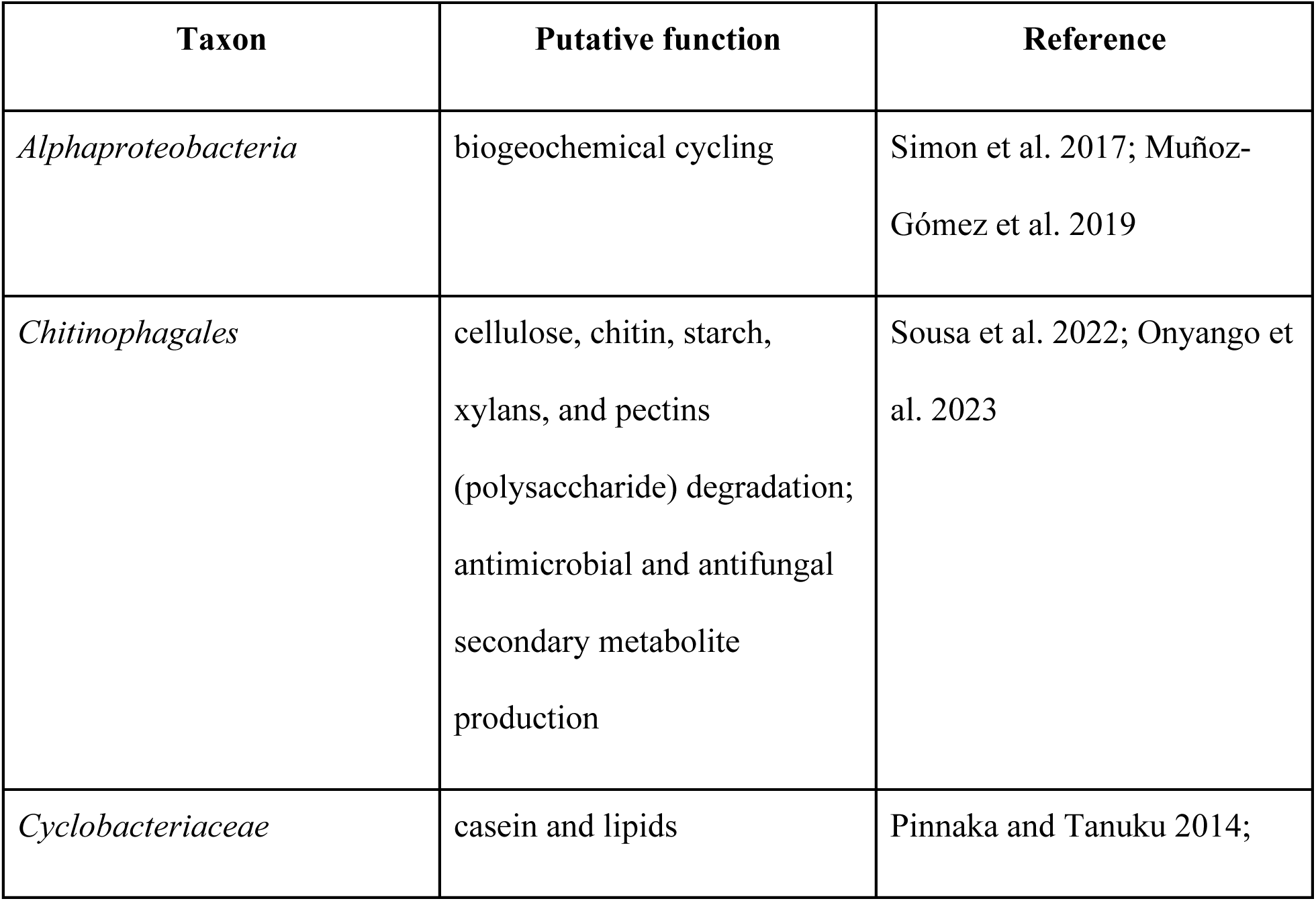
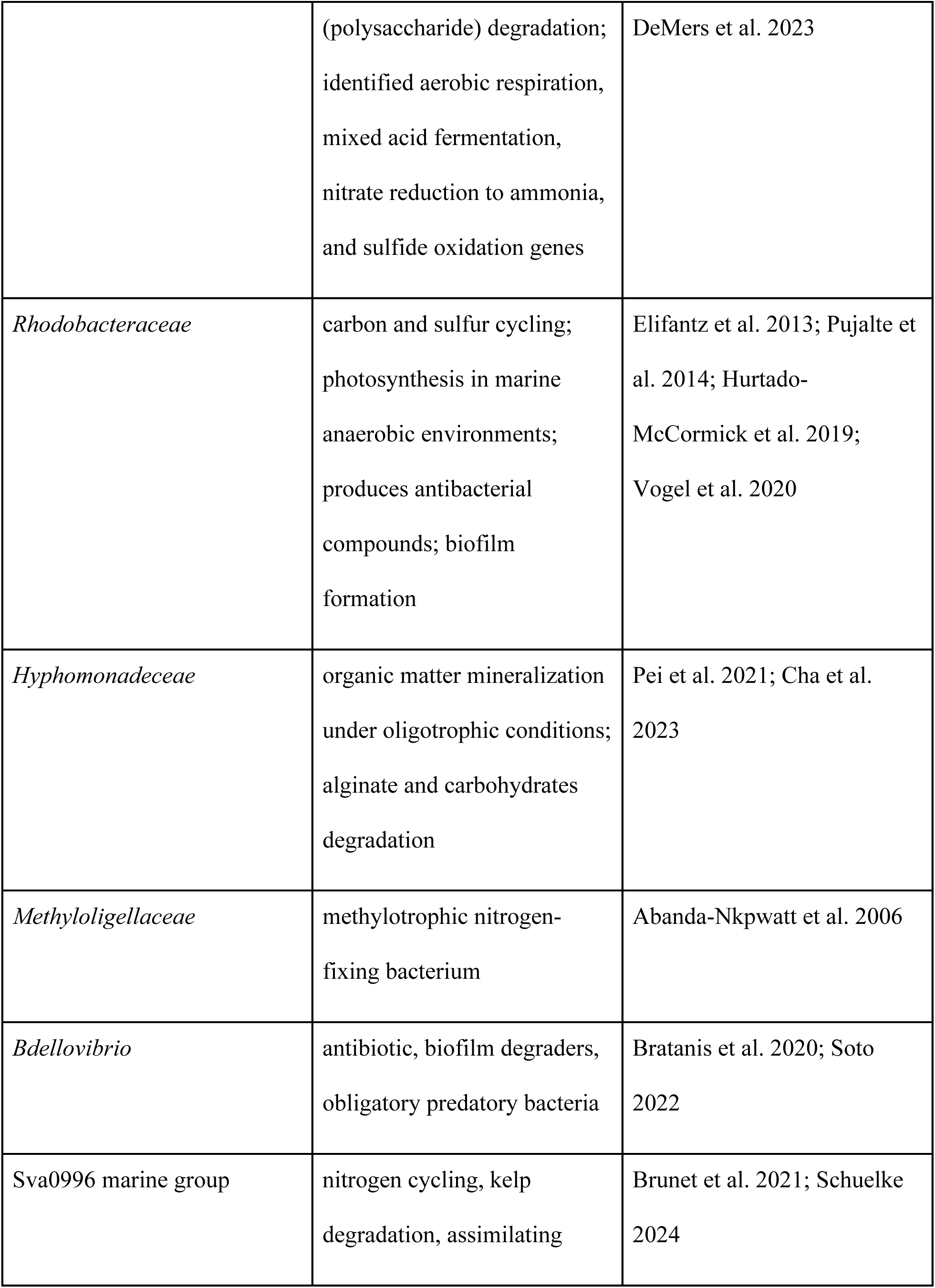
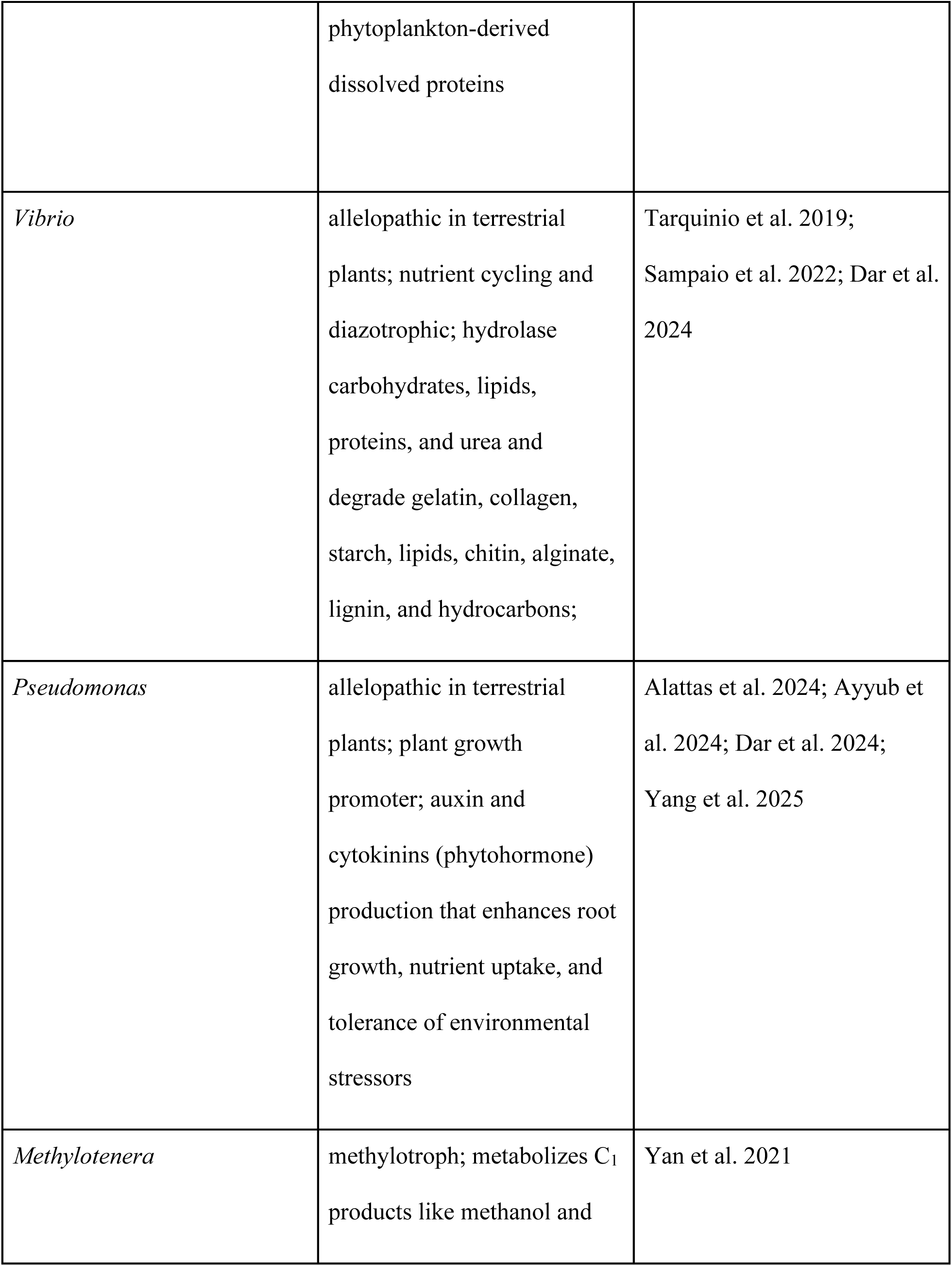
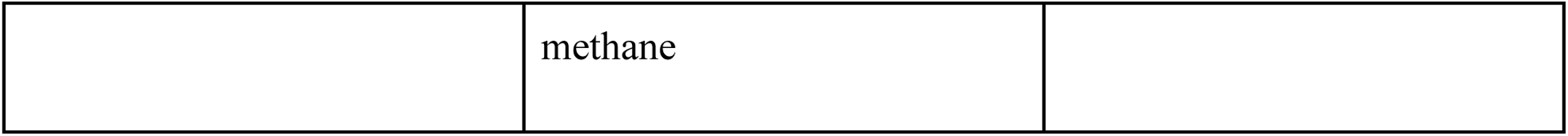
Core microbiome taxa identified in mixed and reduced mixed treatments in the *H. stipulacea* heatmap and their putative function.

**Figure S1A.**
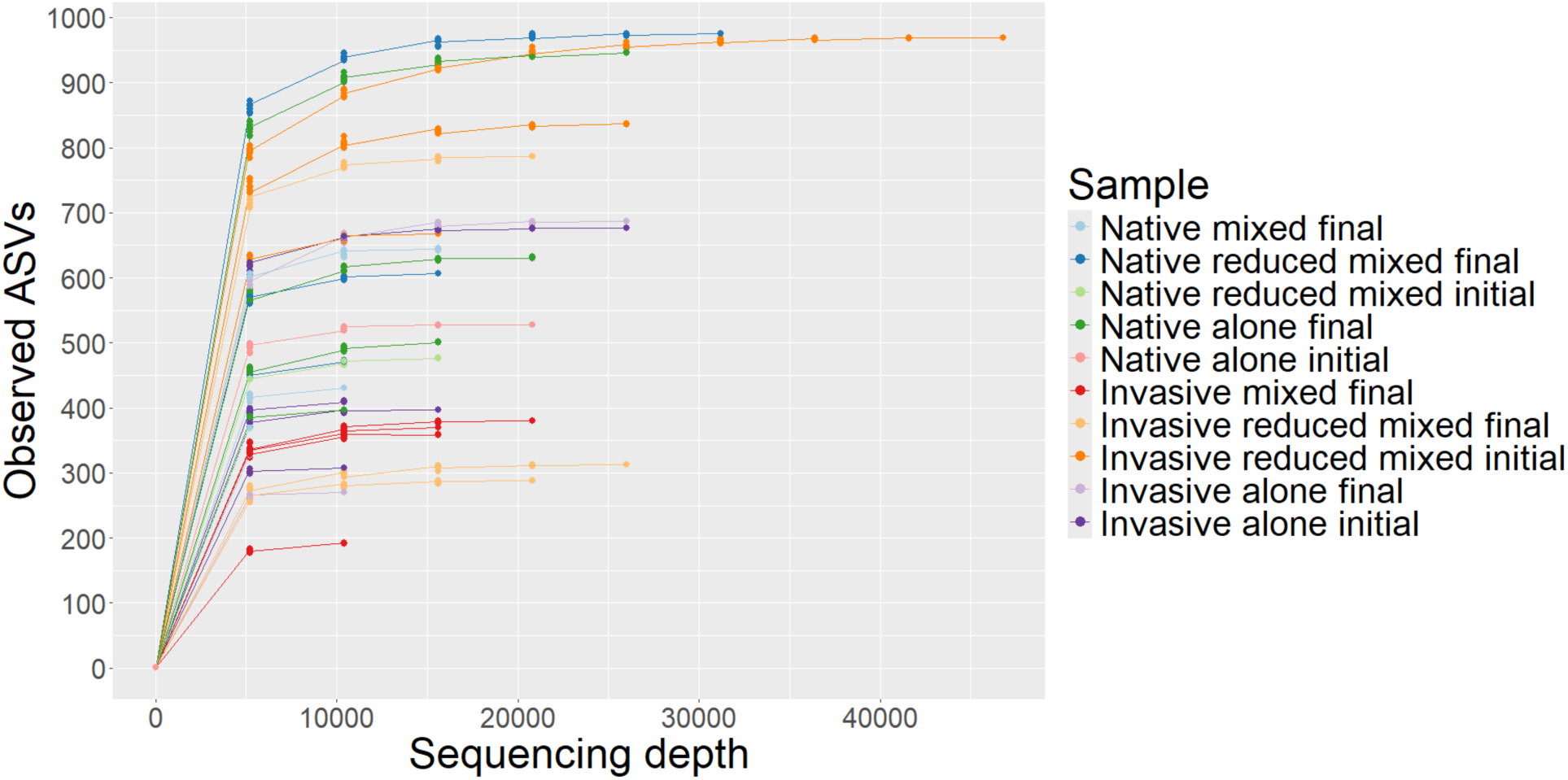
Rarefaction curve of total bacterial diversity for n=40 initial and final samples.

**Figure S1B.**
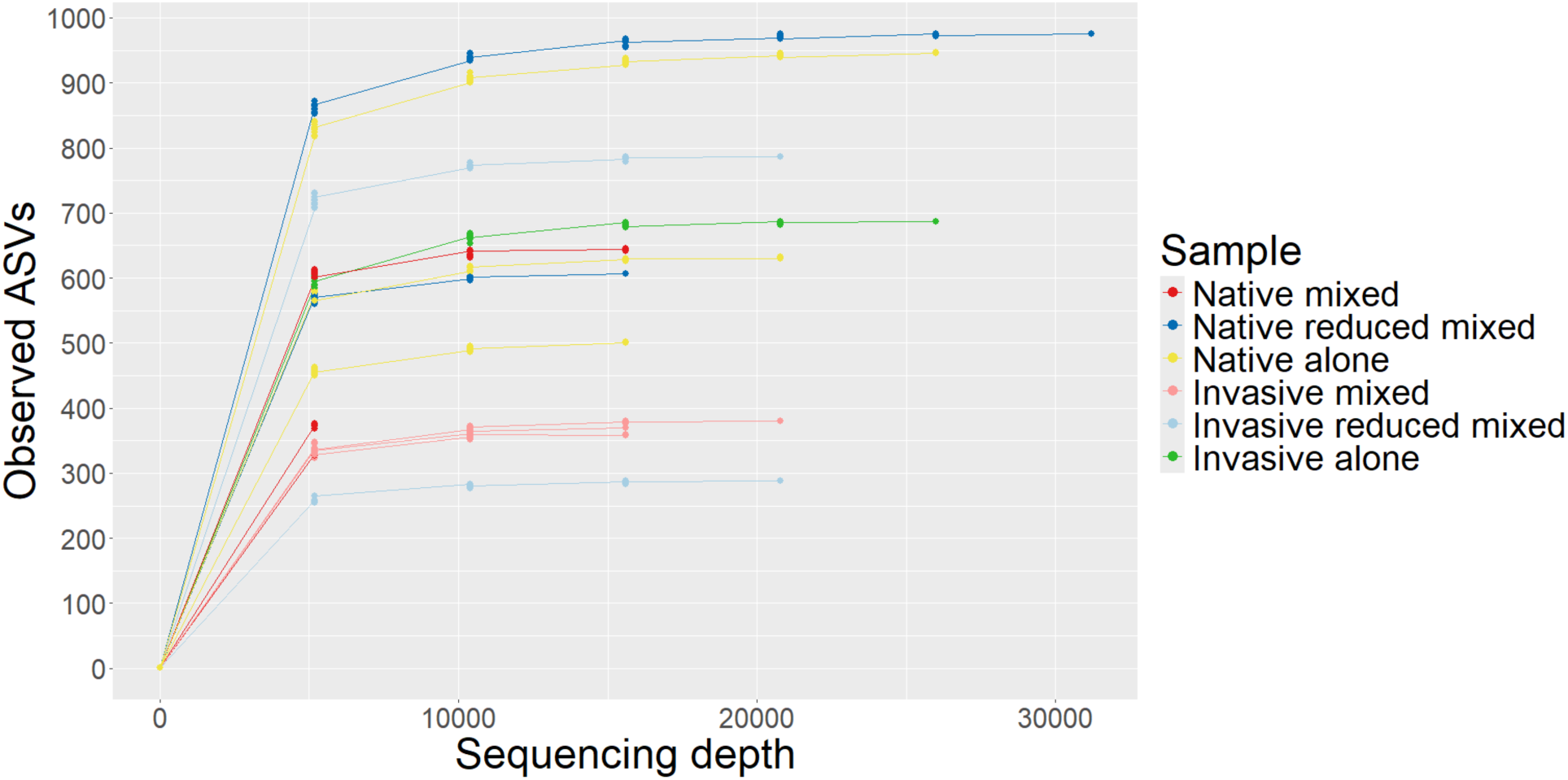
Rarefaction curve of total bacterial diversity for n=18 final samples.

